# Autologous biopsy-derived co-culture platform for interrogation of intestinal epithelial-T cell crosstalk

**DOI:** 10.64898/2026.08.03.742484

**Authors:** Joram Mooiweer, Sajid Anwar, Nathan Ribeiro, Aarón Daniël Ramírez Sánchez, Hanna Simpson, Eline Smits, Renée Anna Maria Moerkens, Jody Gelderloos-Arends, Rutger Modderman, Gieneke Gonera, Margreet Wessels, Cisca Wijmenga, Sebo Withoff, Iris Helene Jonkers

**Affiliations:** Department of Genetics, University Medical Center Groningen, University of Groningen, Groningen, The Netherlands; Department of Genetics, Physical Anthropology and Animal Physiology, University of the Basque Country UPV/EHU, Leioa, Spain; Ombion Centre for Animal-free Biomedical Translation, University of Utrecht, Utrecht, The Netherlands; Department of Pediatrics, Wilhelmina Ziekenhuis Assen, Assen, The Netherlands; Department of Pediatrics, Rijnstate Hospital, Arnhem, The Netherlands

## Abstract

Interactions between intraepithelial lymphocytes (IELs) and the intestinal epithelium are central to mucosal homeostasis and disease. However, mechanistic in vitro studies describing their crosstalk in humans are limited by scarceness of primary material and insufficient knowledge about co-culture requirements. Here, we establish an autologous human duodenal IEL–organoid co-culture system encompassing expandable and bankable IEL and organoid protocols, with co-culture conditions that allow viability of both cell types. This system enables successive interrogation of lympho-epithelial interactions starting from minimal biopsy material. Under baseline conditions, CD45⁺CD8⁺CD103⁺TCRαβ⁺ IELs retain tissue-residency and effector features and induce an epithelial interferon response and chemokine production, without overt epithelial apoptosis. IL-15 and IL-21, essential cytokines involved in IEL-activation in intestinal enteropathies like celiac disease, increases granzyme B expression and interferon-γ secretion but do not trigger epithelial cell death. However, enforcing IEL-epithelial contact using an anti-CD3-anti-Ep-CAM bispecific antibody induces epithelial apoptosis accompanied by increased tumor necrosis factor (TNF) and FAS-ligand (FASLG) secretion. These findings validate the platform’s ability to resolve non-destructive and cytotoxic lympho-epithelial interaction and provide a tractable system for studying intestinal inflammation and immune-mediated epithelial cell death.

## Introduction

Intestinal homeostasis is maintained through continuous interactions between immune cells located at the mucosal barrier, the intestinal epithelia, and the microbiome. Intraepithelial lymphocytes (IELs) play an important role in homeostasis, as they reside in direct contact with epithelial cells, and contribute to barrier surveillance, epithelial repair, and immune-mediated tissue damage ^1^. Because of this intimate spatial relationship, experimental systems that allow for controlled study of human IEL–epithelial interactions are important for understanding mucosal homeostasis and inflammatory disease.

Several populations of IELs contribute to intestinal homeostasis and disease, including conventional TCRαβ⁺ IELs, TCRγδ⁺ IELs, and innate-like lymphocyte populations with distinct regulatory and effector functions^2^. The predominant IEL population in the human small intestine are the CD8⁺CD103⁺TCRαβ⁺ IELs^1^. This subset is of particular pathological relevance in celiac disease (CeD), where it promotes epithelial damage through release of cytotoxic mediators such as granzyme B (GzmB) and IFN-γ and the engagement of NK-like activating receptors ^1,3–5^. Dysregulated IEL responses have also been associated with other chronic inflammatory disorders of the intestine, including inflammatory bowel disease (IBD), where disturbed lymphoepithelial interactions contribute to epithelial injury and barrier dysfunction^6–8^. All these features position CD8⁺CD103⁺TCRαβ⁺ IELs as a particularly relevant population to capture when modeling human lymphoepithelial interactions. However, despite growing recognition of the importance of IELs in intestinal health and disease, human-relevant experimental systems for studying IEL–epithelial interactions remain limited ^9,10^.

Human intestinal organoids provide powerful models for epithelial biology because they preserve key properties of primary intestinal tissue while also offering experimental accessibility ^11–14^. However, conventional organoid systems are epithelial cell−only and lack the immune compartment central to epithelial regulation *in vivo*. Immune-organoid co-culture models would address this caveat, e.g. through incorporation of circulating or tissue-resident immune populations ^9,10^. Most current IEL–organoid co-culture systems rely primarily on murine cells ^15–22^, underscoring the difficulty of establishing comparable human models. A likely explanation for this is the limited availability of human biopsy material and the low yield of the primary IELs that can be recovered for downstream experimentation. Furthermore, most human IEL– organoid co-culture systems lack donor-matched immune compartments, limiting our ability to capture inter-individual heterogeneity and examine patient-specific immune and epithelial interactions. Recent studies have begun to address this challenge by establishing autologous intestinal immune-organoid systems that retain or reconstitute tissue-resident immune compartments ^23–26^. However, we still lack a defined and renewable human-based immune-organoid platform tailored for the controlled study of human IEL–epithelial interactions.

Here, we establish a platform for IEL–epithelial cultures using donor-matched cells derived from duodenal biopsies. We developed a workflow to isolate, expand, bank, and re-activate human CD45⁺CD8⁺CD103⁺TCRαβ⁺ IELs from minimal starting material while also generating matched intestinal organoids from the same donors. Crucially, we define a consensus co-culture medium that sustains both epithelial organoids and IELs. These features overcome several key technical barriers that have limited the use of primary human IELs in organoid-based assays. In short, this paper presents a technically defined, autologous, and renewable human IEL–organoid co-culture platform.

## Results

### Collection of intestinal biopsies for isolation of adult stem cells and IELs

To establish immune-competent organoid co-cultures with short-lived tissue-derived IEL cell lines, we collected four duodenal pinch biopsies per donor. These were then cut into small fragments and cryopreserved following protocols previously established by our group ^27^. This approach allows for long-term storage and revival of cells, thereby decoupling downstream processing from tissue acquisition and avoiding the time-pressure constraints of handling fresh samples.

### Isolation, expansion, and banking of tissue-resident IELs

Upon thawing, two pinch biopsies per individual were dissociated into single-cell suspensions, and CD45⁺CD8⁺CD103⁺TCRαβ⁺ T cells were isolated by FACS (Fig. 1a,b). Expression of CD103, an integrin that interacts with epithelial E-cadherin, was used to enrich T cells with tissue-resident and intraepithelial properties. For simplicity, these CD103⁺ T cells are hereafter referred to as IELs. Sorting yielded IEL numbers ranging from 200 to 25,000 (Fig. 1c). These cells were then expanded on irradiated feeder cells in the presence of IL-2 and PHA-L, yielding between 13 million and 106 million cells, respectively, after 21 days (Supplementary Fig. 1a and Fig. 1c), which were cryopreserved in batches of 750,000 cells per vial. Expanded IEL cultures showed no detectable feeder carryover and consisted exclusively of CD8+TCRαβ+ IELs, confirmed by flow cytometry (Fig. 1d). However, CD103 expression was reduced (7.7–36.6%, Fig. 1d) compared to the 100% expression on freshly isolated IELs (Fig. 1b), possibly due to the absence of epithelial cell−derived signaling in the feeder mix (Fig. 1d). For experimental use, cryopreserved IELs were re-expanded, yielding between 23 million and 49 million cells after 10 days (Fig. 1e). To assess whether the *in vitro* expansion and freezing cycle altered key phenotypic features, we evaluated the exhaustion markers PD-1 and TIGIT and the senescence marker CD57, but none of these markers were detectably present on the cell surface (Supplementary Fig. 1e−g). In addition, the frequency of cells positive for the effector molecule GzmB and the gut homing marker CCR9 remained high (Supplementary Fig. 1c,d). To promote re-expression of the tissue-residency marker CD103, the expanding cultures were supplemented with TGF-β on day 9, 24 hours prior to conducting functional assays ^28^, resulting in an increase in CD103 on the cell surface (Fig. 1f) without altering the levels of GzmB, CCR9 (Supplementary Fig. 1c,d,h), or exhaustion- and senescence-associated markers (Supplementary Fig. 1e,f). Altogether, this approach robustly and reproducibly enables expansion, preservation, and re-expansion of human IELs expressing tissue-residency markers from endoscopic pinch biopsies.

**Fig. 1.**
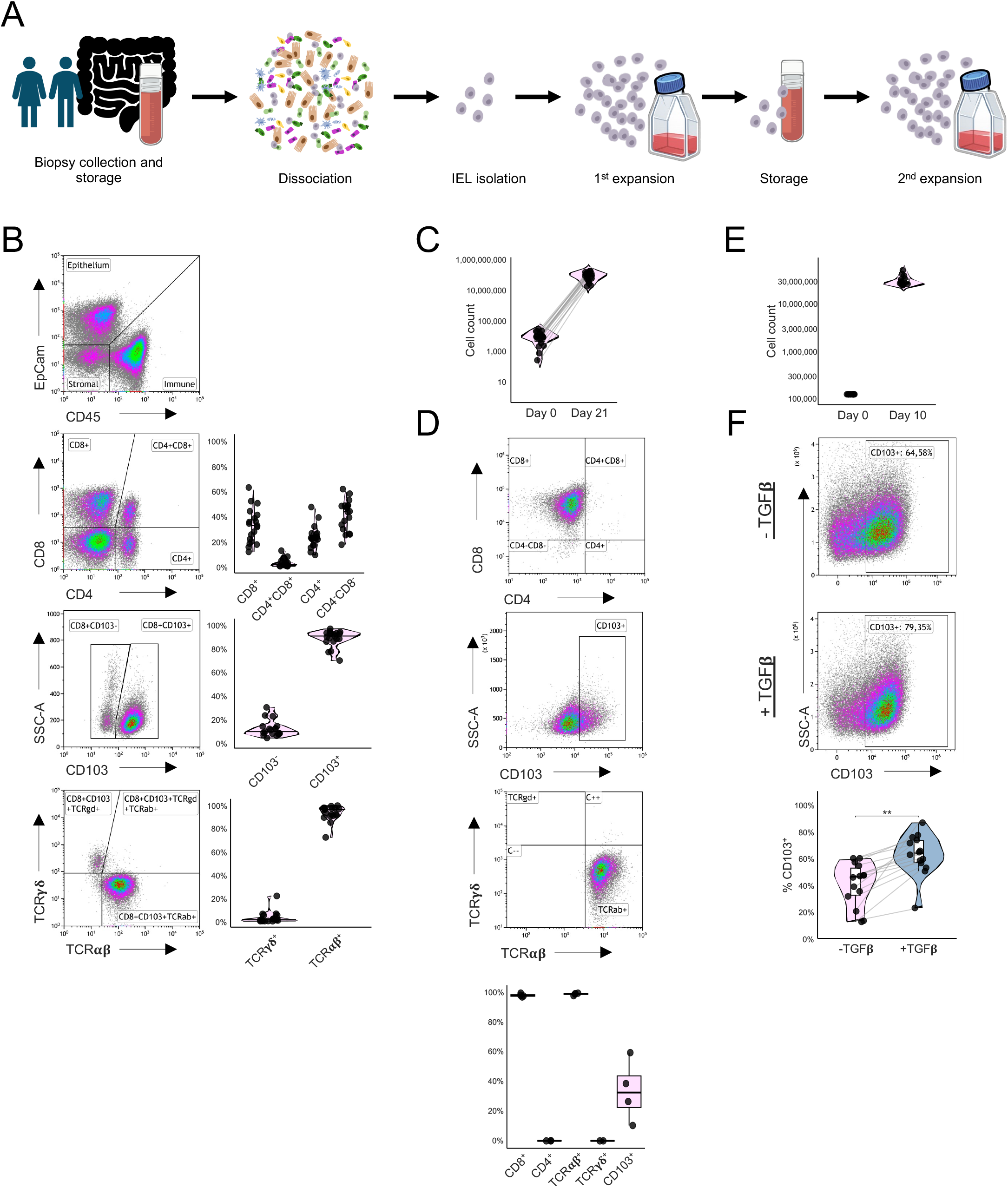
Biopsy-derived intestinal IELs can be expanded, banked and re-expanded while retaining key phenotypic features. **a** Schematic overview of the workflow used to isolate, expand, cryopreserve, and re-expand short-lived intraepithelial lymphocyte (IEL) lines from intestinal biopsies. **b** Representative flow cytometry plots of dissociated intestinal biopsies and quantification of T cell populations. IELs were defined as CD45⁺CD8⁺CD103⁺TCRαβ⁺ cells (n = 19 donors). **c** IEL cell counts after isolation (day 0) and after the first expansion round (day 21) (n = 19 donors). **d** Representative flow cytometry plots and quantification of IEL phenotype after the first expansion round (n = 4 donors). **e** IEL cell counts after revival of cryopreserved IELs (day 0) and after re-expansion (day 10) (n = 15 IEL lines). **f** Representative flow cytometry plots and quantification of CD103 expression in re-expanded IELs cultured with or without TGF-β stimulation (10 ng/mL for 24 h) (n = 14 IEL lines). Each dot represents one donor or IEL line. In (**f**), statistical significance was assessed using paired t-tests with Benjamini-Hochberg correction for multiple comparisons where indicated. *P < 0.05, **P < 0.01, ***P < 0.001, ****P < 0.0001.

### Generation of adult stem cell−derived organoids

In parallel with establishing the IEL lines, duodenal organoids were generated from two additional donor-matched pinch biopsies. Following biopsy fragmentation and embedding in Matrigel domes, the organoids expanded progressively over approximately 3 weeks, forming robust, fast-growing cultures suitable for cryopreservation (Supplementary Fig. 1i). ^29^Organoid cultures were successfully established for all donors to facilitate personalized autologous co-culture systems.

### Development of a co-culture medium to sustain both organoids and IELs

To establish IEL–organoid co-cultures, we first assessed whether organoids and IELs could be maintained under the standard culture conditions for the other.

To enable reliable quantitative assessment of organoid growth in an ‘open-top’ configuration compatible with downstream co-culture experiments ^29^ organoids were cultured on 2D Matrigel beds rather than in Matrigel domes. Organoid formation on the Matrigel beds was robust, but they collapsed within 24 hours after switching to IEL medium (Fig. 2a,b) and did not recover at later time points (Supplementary Fig. 2a,b). Organoid collapse was also observed when they were cultured in standard Matrigel domes with IEL medium (Supplementary Fig. 2c and Supplementary Video 1), indicating that the loss of organoid integrity was driven by medium composition rather than culture geometry.^29,29^

**Fig. 2.**
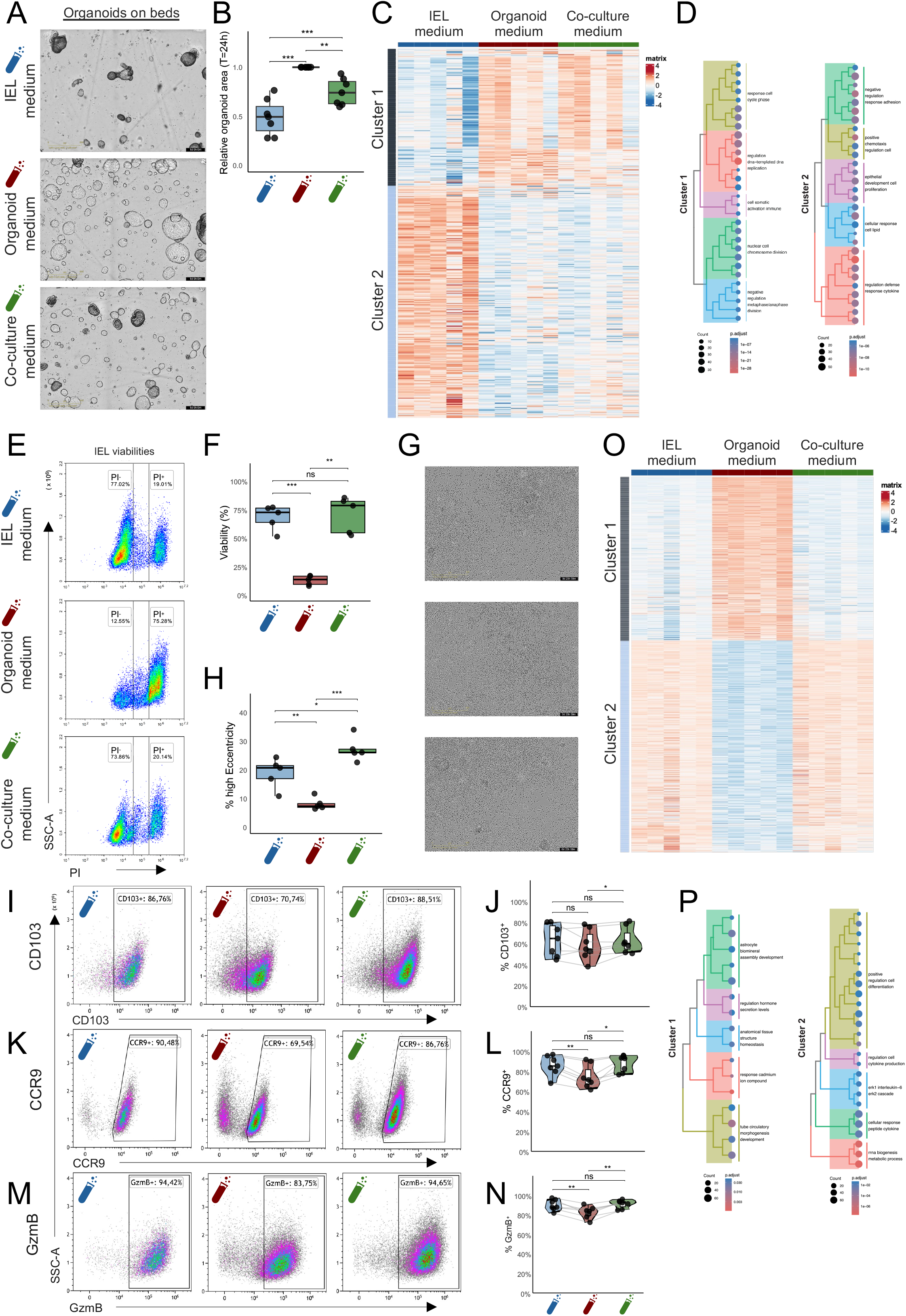
A shared co-culture medium sustains intestinal organoids and IELs. **a** Representative brightfield images of intestinal organoids grown on Matrigel beds after switching to IEL medium, organoid medium, or co-culture medium. **b** Quantification of relative organoid surface area 24 h after medium switch. Values are normalized to organoids maintained in organoid medium, which was set to 1. **c** Heatmap showing transcriptional profiles of organoids cultured for 6 h in the media indicated. Genes were grouped into two main clusters based on the expression patterns based on differentially expressed genes across conditions. **d** Over-represented pathways associated with the organoid gene clusters defined in C. **e** Representative propidium iodide (PI) live/dead staining of IELs 24 h after switching to the indicated media. **f** Quantification of IEL viability based on PI staining. Each dot represents one IEL line (n = 5). **g** Representative phase-contrast images of IELs 24 h after switching to the indicated media. **h** Quantification of IELs with high eccentricity 24 h after medium switch. Each dot represents one IEL line (n = 5). (**i**–**n**) Representative flow cytometry plots and quantification of CD103⁺, CCR9⁺, and GzmB⁺ IEL populations after culture in the media indicated for 12 h (n = 7 IEL lines). **o** Heatmap showing transcriptional profiles of IELs cultured for 6 h in the media indicated. Genes were grouped into two main clusters based on expression patterns across conditions. **p** Over-represented pathways associated with the IEL gene clusters defined in o. In all quantification plots, each dot represents one organoid line or IEL line, as indicated. Statistical significance was assessed using paired t-tests with Benjamini-Hochberg correction for multiple comparisons where indicated. *P < 0.05, **P < 0.01, ***P < 0.001, ****P < 0.0001.

Similarly, IELs did not survive in standard organoid medium (Fig. 2e,f). Therefore, to develop a general co-culture medium, we deleted Prostaglandin E2 ^30,31^, nicotinamide ^32^, TGF-β inhibitor A83-01 ^28^, and MAPK inhibitor SB202190 ^33–35^ from the standard organoid growth media because they have been reported to impair T cell growth, activity, or tissue-residency phenotypes. The medium was also supplemented with the T cell– supporting cytokines IL-2 and IL-7 ^16,17,23–26,29^ to promote IEL survival and activation.

Organoids cultured in the co-culture medium remained morphologically similar to those cultured in standard organoid medium (Fig. 2a and Supplementary Video 1), although their surface area in co-culture media was decreased relative to organoids cultured in standard organoid medium (Fig. 2b). Transcriptomic profiling of organoids exposed to IEL medium revealed a distinct RNA profile (Fig. 2c and Supplementary Tables 1 and 2) with genes involved in inflammatory pathways being upregulated relative to organoids grown in co-culture or standard organoid medium (Fig. 2e and Supplementary Fig. 2b), indicative of increased cellular stress, dysbiosis and damage. In contrast, organoids cultured in co-culture medium closely resembled organoids maintained in standard organoid medium (Fig. 2c) and displayed increased expression of genes involved in DNA replication pathways (Fig. 2d, Supplementary Fig. 2e and Supplementary Table 3), indicative of proliferation and homeostasis.

IELs also remained viable in co-culture medium (Fig. 2e,f) and maintained their characteristic eccentric morphology as in standard IEL medium (Fig. 2g, h and Supplementary Fig. 2d). Flow cytometric analysis further showed that the frequency of cells expressing the IEL-associated markers CD103 (Fig. 2i,j), CCR9 (Fig. 2k,l), and GzmB (Fig. 2m,n) was maintained in co-culture medium at levels comparable to those in standard IEL medium. Likewise, GzmB median fluorescent intensity (MFI) remained unchanged (Supplementary Fig. 2f), indicating that IELs’ cytotoxic potential was maintained. In contrast, culturing IELs in standard organoid medium caused a reduction in the expression of GzmB (Fig. 2m,n), and CCR9 (Fig. 2k,l). Moreover, when cultured in organoid medium, IELs presented a distinct transcriptional group (Fig. 2o and Supplementary Tables 4 and 5) characterized by enrichment of gene sets associated with ion-response and broader tissue homeostasis/development, indicative of a medium-driven adaptation or stress response. In contrast, IELs cultured in co-culture medium resembled IELs maintained in standard IEL medium (Fig. 2o) and were enriched for pathways related to rRNA processing and cytokine-responsive signaling, consistent with preservation of biosynthetic activity and general immune cell fitness (Fig. 2p and Supplementary Table 6).

Together, these results demonstrate that the consensus co-culture medium sustains both the epithelial and lymphocyte compartments in physiologically relevant states.

### Co-culturing IELs and intestinal organoids

Having confirmed the functionality of the co-culture medium, we aimed to establish stable and reproducible co-cultures of IELs and organoids. Initially, we observed that organoids embedded in Matrigel domes did not permit efficient IEL–epithelium interactions, with IELs accumulating in the dome periphery and only rarely penetrating the Matrigel matrix (Supplementary Fig. 3a and Supplementary Video 2). We therefore adopted the Matrigel-bed format ^29^, which not only supported robust organoid growth in co-culture medium (Fig. 2a,b) it also allowed IELs to be added directly onto the organoid cultures, promoting close spatial association between the two compartments (Fig. 3a).

**Fig. 3.**
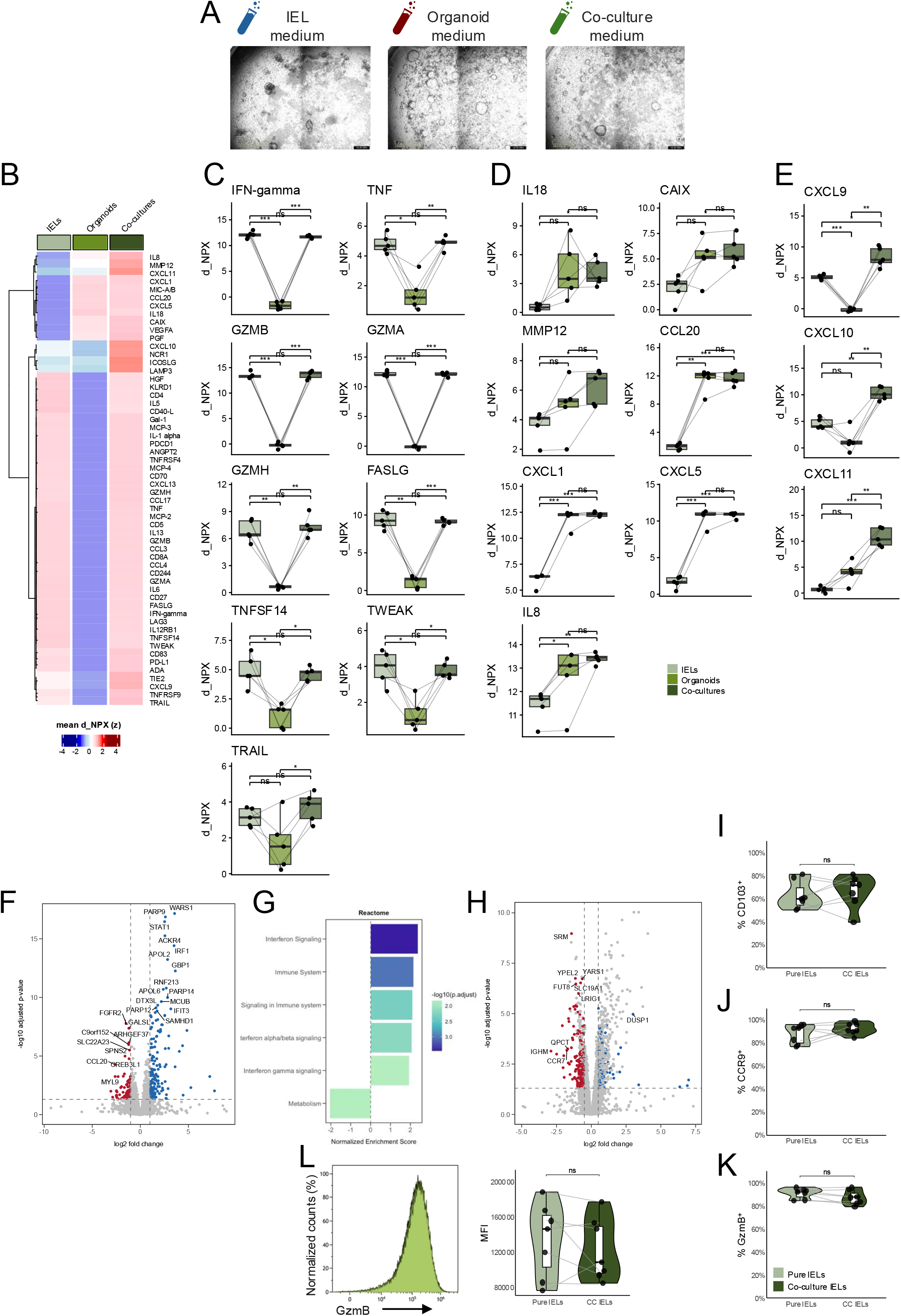
IEL–organoid co-culture induces epithelial interferon signalling with limited changes in IEL state. **a** Representative images of IEL–organoid co-cultures established on Matrigel beds and cultured for 24 h in IEL medium, organoid medium, or co-culture medium. **b** Heatmap of Olink-detected cytokines, chemokines, and immune-mediators in conditioned media from IEL monocultures, organoid monocultures, and IEL–organoid co-cultures grown in co-culture medium (n = 5). Values are shown as row-wise z-scores of mean blank-corrected NPX values. **c** Quantification of selected IEL-associated effector and cytotoxic mediators detected by Olink in conditioned media. **d** Quantification of selected organoid-associated chemokines and epithelial mediators detected by Olink in conditioned media. **e** Quantification of co-culture-induced interferon-associated chemokines detected by Olink in conditioned media. In C–E, each dot represents one conditioned media sample (n = 5). **f** Transcriptomic analysis of organoids cultured alone or recovered from IEL–organoid co-cultures after 6 h, as determined by bulk RNA-seq (n = 5). **g** Pathway analysis of transcriptional changes in organoids after co-culture. **h** Transcriptomic analysis of IELs cultured alone or recovered from IEL–organoid co-cultures after 6 h, as determined by bulk RNA-seq (n = 5). (**i**–**k**) Quantification of CD103⁺, GzmB⁺, and CCR9⁺ IEL populations in IEL monocultures and after recovery from co-cultures. **l** Quantification of intracellular GzmB levels in GzmB⁺ IELs, shown as median fluorescence intensity (MFI). In I–L, each dot represents one individual IEL line (n = 7). Statistical significance was assessed using paired tests with Benjamini-Hochberg correction for multiple comparisons where indicated. *P < 0.05, **P < 0.01, ***P < 0.001, ****P < 0.0001.

To visualize IEL behavior in co-culture, IELs were pre-stained with a red fluorescent dye and imaged by time-lapse microscopy over 20 hours (Supplementary Video 3). In organoid medium, IELs were largely static, whereas IELs in IEL- or co-culture media were very motile. IELs also seemed to interact more with organoids in co-culture medium, as transient red signal was detected at the organoid periphery. Consistent with this, Recaldin et al. previously reported that tissue-derived T cells in immune-organoid cultures are highly migratory and move dynamically within both the extracellular medium (ECM) and epithelial layers, thereby promoting repeated epithelial contacts ^24^. Matrigel-bed co-cultures therefore enable dynamic IEL behavior and physical proximity to organoids.

### IEL–organoid co-cultures induce an epithelial interferon response

To characterize molecular crosstalk between IELs and intestinal organoids, we profiled cytokine and chemokine secretion in IEL monocultures, organoid monocultures, and IEL–organoid co-cultures, all grown in co-culture medium (Fig. 3b–e and Supplementary Table 7 and 8). Most of the proteins detected in the co-culture supernatants overlapped with factors present in either IEL- or organoid-monoculture supernatants, indicating that most signals can be attributed to one of the two compartments (Fig. 3b and Supplementary Fig. 3). Classical IEL effector molecules such as IFNγ, TNF, GzmB, GzmA, and GzmH were abundant in IEL monocultures and showed comparable levels in co-cultures (Fig. 3c). In addition, in both mono- and co-cultures, IELs secreted multiple cytotoxic and co-stimulatory mediators, including FASLG, TNFSF14 (LIGHT), TWEAK, and TRAIL, consistent with a pre-activated cytotoxic phenotype. The IEL-origin of these factors was confirmed on the transcriptomic level, which showed expression of these genes only in IELs and not in organoids (Supplementary Fig. 3d). Cytokine production upon co-culture was, however, not increased, suggesting that the IELs are in a primed-effector state independent of epithelial interaction (Fig. 3c).

Organoid monocultures were characterized by secretion of epithelial-derived chemokines and stress-associated factors, including IL-18, CCL20, CXCL1, CXCL5, and IL-8 (Fig. 3d), which remained unchanged in co-culture supernatants. The organoid origin of these factors was confirmed by the RNA expression levels of these genes (Supplementary Fig. 3e). This shows that organoids are already secreting these immunogenic proteins at baseline, and this is not further enhanced by the presence of IELs.

Notably, a subset of proteins displayed synergistic induction under co-culture conditions. Specifically, the interferon-inducible chemokines CXCL9, CXCL10, and CXCL11 were detected at lower levels in monocultures but were highly secreted (Fig. 3e) and expressed (Supplementary Fig. 3f) by organoids in co-cultures, either due to direct IEL–epithelial interactions or IEL-derived IFNγ (Fig. 3c). Consistent with these observations, transcriptomic analysis of organoids revealed robust induction of interferon-response genes following co-culture with IELs (Fig. 3f and Supplementary Table 9). Gene set enrichment analysis confirmed activation of type I and type II interferon signaling and cytokine-mediated immune responses (Fig. 3g and Supplementary Table 10). In contrast, IELs exhibited minimal transcriptional changes between the monoculture and co-culture conditions (Fig. 3h and Supplementary Table 11), in line with the largely unchanged secretion profile we observed at the protein level (Fig. 3b,c). Cell surface marker expression also showed no major changes in CD103⁺, CCR9⁺, and GzmB^±^ populations (Fig. 3i−k) or in intracellular GzmB levels (Fig. 3l).

Taken together, these data indicate that IELs maintain a stable, pre-activated effector state when cultured with organoids, but they can still induce a robust interferon response in the epithelium through secretion of IFNγ. This interaction recapitulates a key feature of mucosal immunity in which tissue-resident IELs promote epithelial defense programs without inducing overt cytotoxicity or cell death.

### IL-15 and IL-21 potentiate IEL effector features

While the co-culture medium containing IL-2 and IL-7 already supported a basal effector-like state, as reflected by the presence of inflammatory mediators in the culture system, we next asked whether a more strongly pro-inflammatory intestinal milieu could further enhance IEL activation and promote apoptosis of epithelial cells.

First, we investigated if a pro-inflammatory milieu affects IELs and organoids in monocultures. IELs and organoids were exposed to IL-15 and IL-21, two cytokines known to enhance IEL effector function and intestinal inflammation ^4,36^. Cultures were also treated with the bispecific antibody BIS1, a clinical-grade CD3×EpCAM bispecific antibody^37,38^ that targets T cells to EpCAM^+^ epithelial cells. Organoids maintained in co-culture medium with or without IL-15 and IL-21 or BIS1 exhibited comparable growth rates (Fig. 4a,b and Supplementary fig. 4a) and no transcriptomic variation (Supplementary Fig. 4b and Supplementary Table 12), indicating that neither pro-inflammatory cytokines nor BIS1 affect epithelial cell growth and behavior.

**Fig. 4.**
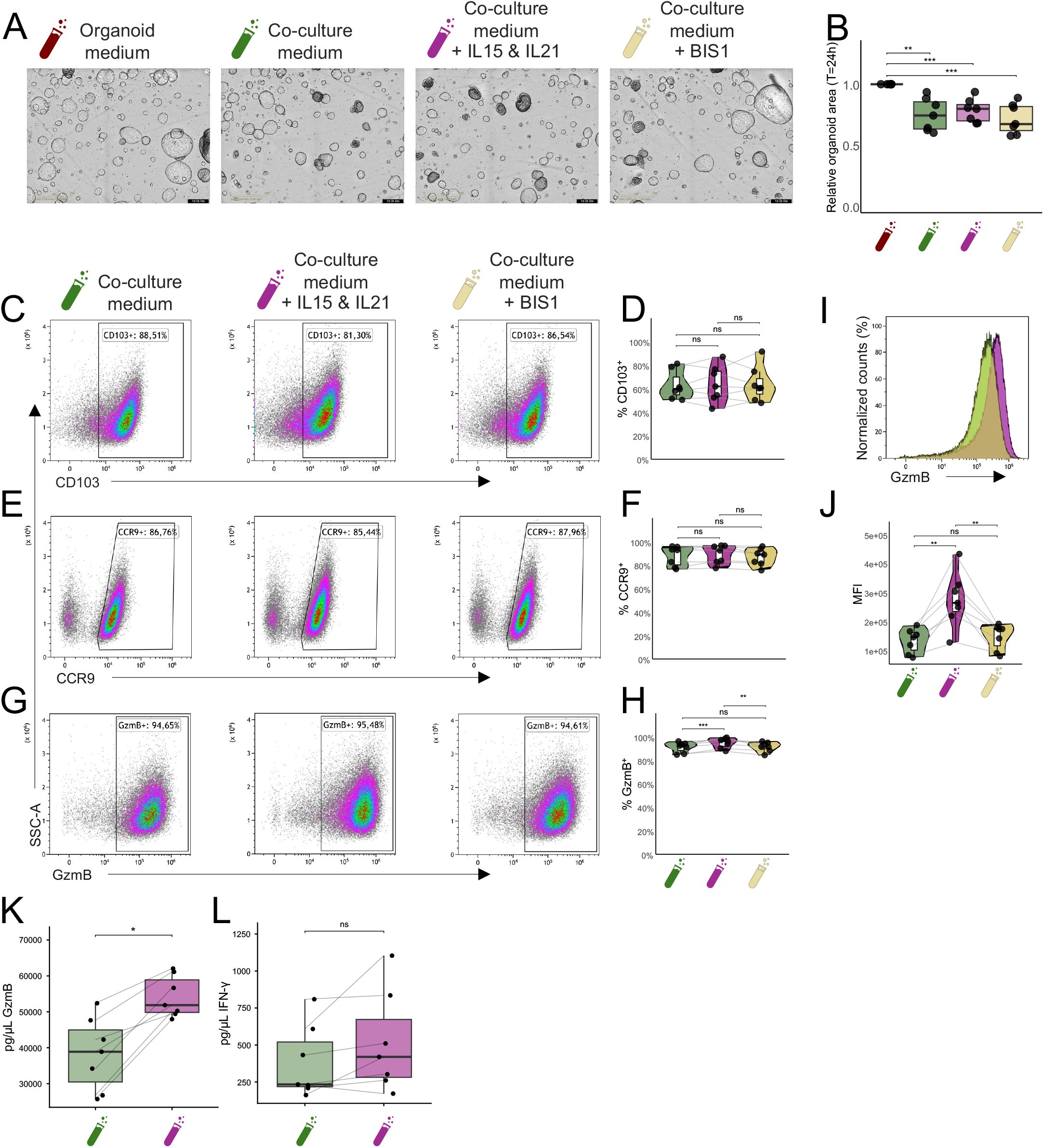
IL-15 and IL-21 enhance IEL effector features without affecting organoid growth. **a** Representative brightfield images of intestinal organoids grown on Matrigel beds 24 h after switching to organoid medium, co-culture medium, co-culture medium supplemented with IL-15 and IL-21, or co-culture medium supplemented with BIS1. **b** Quantification of relative organoid surface area 24 h after medium switch. Values were normalized to organoids maintained in organoid medium, which was set to 1 (n = 7). **c**, **e**, **g** Representative flow cytometry plots of CD103⁺, CCR9⁺, and GzmB⁺ IEL populations cultured in co-culture medium, co-culture medium supplemented with IL-15 and IL-21, or co-culture medium supplemented with BIS1. **d**, **f**, **h** Quantification of CD103⁺, CCR9⁺, and GzmB⁺ IEL populations under the indicated conditions (n = 7). **i** Representative histogram of intracellular GzmB expression in IELs cultured under the indicated conditions. **j** Quantification of intracellular GzmB levels in GzmB⁺ IELs, shown as median fluorescence intensity (MFI) (n = 7). **k**, **l** Concentrations of GzmB (k) and IFN-γ (**l**) determined by ELISA on culture supernatant of IEL monocultures cultured in co-culture medium or co-culture medium supplemented with IL-15 and IL-21 (n = 7). For quantification plots, each dot represents one organoid line or IEL line, as indicated. Statistical significance was assessed using paired t-tests with Benjamini-Hochberg correction for multiple comparisons where indicated. *P < 0.05, **P < 0.01, ***P < 0.001, ****P < 0.0001.

Addition of the pro-inflammatory cytokines IL-15 and IL-21 to IEL monocultures had only a modest effect on gene expression, with 14 genes differentially expressed, including immune-regulatory factors such as IRF4 and SOCS3 (Supplementary Fig. 4c and Supplementary Table 13). CD103 and CCR9 levels remained consistently unchanged upon IL-15 and IL-21 stimulation (Fig. 4c−f). Although GzmB expression was already high under baseline conditions, IL-15 and IL-21 stimulation induced a modest but significant increase in GzmB⁺ cell frequency and intracellular GzmB protein levels (Fig. 4g−j), along with elevated GzmB levels in the supernatant (Fig. 4k). This indicates potentiation of IEL effector function. IFNγ levels were, however, unchanged upon IL-15 and IL-21 stimulation of IEL monocultures (Fig. 4l). In addition, BIS1 addition did not affect the cell-surface markers CD103 and CCR9 or intracellular levels of GzmB in IEL monocultures (Fig. 4c−j).

Taken together, these findings show that IL-15 and IL-21 have a moderate additive effect on IEL effector function compared to the basic co-culture medium when cultured in isolation, likely because IL-2 and IL-7 already induce a primed-effector state.

### Pro-inflammatory milieu alone is insufficient to drive IEL-mediated epithelial cell death

We next asked whether IEL effector potential translates into functional cytotoxicity directed towards epithelial cells. To quantify IEL-mediated epithelial killing, we developed a live-cell apoptosis assay using a fluorescent caspase tracker compatible with IncuCyte time-lapse imaging. IELs and organoids cultured separately served as negative controls to assess spontaneous apoptosis, while puromycin treatment was used as a positive control defining maximal caspase activation (Supplementary fig. 5A). Here we observed a strong caspase signal in puromycin-treated wells and minimal to no signal in IEL and organoids in monoculture with co-culture medium, under all conditions (Supplementary Fig. 5a–f). Puromycin-caused maximal cell death was used as an internal caspase tracker normalization for subsequent analyses in co-culture experiments.

Co-cultures supplemented with IL-15 and IL-21 did not display increased cell death compared to co-cultures in standard co-culture media. This suggests that, while IL-15 and IL-21 do potentiate cytotoxic capacity of IELs in monocultures (Fig. 4c−j), additional factors may be necessary to induce IEL-mediated killing of epithelial cells in co-cultures (Fig. 5a–c). In contrast, BIS1-treated co-cultures showed extensive apoptosis (Fig. 5a−c). These results confirm that the IEL–organoid system functionally responds to cytokine-mediated and bispecific antibody−mediated T cell engagement and can quantitatively capture T cell–mediated epithelial killing.

**Fig. 5.**
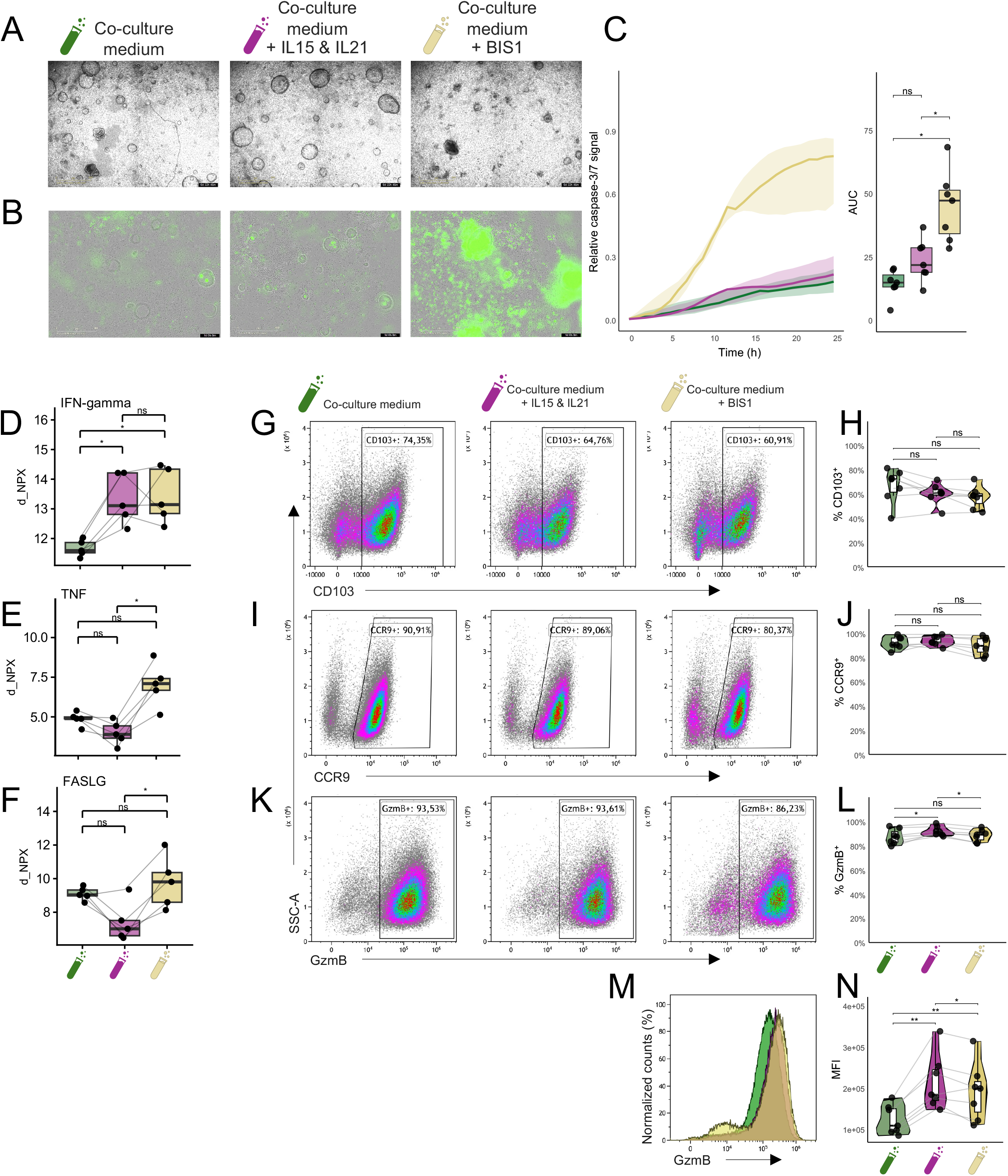
Enforced CD3×EpCAM engagement, but not cytokine priming, induces epithelial apoptosis. **a,b** (**a**) Representative brightfield images and (**b**) phase-contrast images with caspase-3/7 fluorescence of IEL–organoid co-cultures grown in co-culture medium, co-culture medium supplemented with IL-15 and IL-21, or co-culture medium supplemented with BIS1. **c** Quantification of relative caspase-3/7 signal over time in IEL–organoid co-cultures under the indicated conditions. Caspase-3/7 signal was normalized to the maximal signal induced by puromycin-treated positive controls, which was set to 1. Data represent medians ± IQRs. Right panel shows the area under the curve (AUC) of the caspase-3/7 signal shown in the left panel (n = 7). **d**–**f** Olink measurements of IFN-γ, TNF, and FASLG levels in conditioned media from IEL–organoid co-cultures under the indicated conditions, shown as blank-corrected d_NPX values (n = 5). **g**, **i**, **k** Representative flow cytometry plots of CD103⁺, CCR9⁺, and GzmB⁺ IEL populations recovered from co-cultures. **h**, **j**, **l** Quantification of CD103⁺, CCR9⁺, and GzmB⁺ IEL populations under the indicated conditions. **m** Representative histogram of intracellular GzmB expression in IELs recovered from co-cultures. **n** Quantification of intracellular GzmB levels in GzmB⁺ IELs, shown as median fluorescence intensity (MFI). For quantification plots, each dot represents one donor-matched IEL–organoid co-culture or IEL line, as indicated. Statistical significance for caspase-3/7 AUC quantification in C was assessed using paired Wilcoxon tests with Benjamini-Hochberg correction for multiple comparisons. For other quantification plots, statistical significance was assessed using paired tests with Benjamini-Hochberg correction for multiple comparisons where indicated. *P < 0.05, **P < 0.01, ***P < 0.001, ****P < 0.0001.

To identify factors contributing to IEL-mediated epithelial cell death after BIS1 addition, we profiled cytokine and effector molecule secretion in co-culture supernatants (Supplementary Fig. 6 and Supplementary tables 14 and 15). IFN-γ levels were significantly increased upon both IL-15 and IL-21 stimulation and upon addition of BIS1 (Fig. 5d). In contrast, TNF and FASLG were not increased by IL-15 and IL-21, instead showing a modest, non-significant reduction. These observations appear consistent with prior studies in human IELs indicating that IL-15 and IL-21 preferentially enhance perforin- and granzyme-associated effector programs rather than FasL- or TNF-mediated cytotoxicity ^36,39,40^. Further analysis of cell-surface markers showed no changes in CD103^+^ and CCR9^+^ cells upon IL-15 and IL-21 stimulation and BIS1 treatment (Fig. 5g−j). GzmB intracellular levels were increased upon IL-15 and IL-21 stimulation but not upon BIS1 treatment, indicating that BIS1-mediated cell death is not dependent on GzmB (Fig. 5k−n). Moreover, increased IFN-γ levels in the co-cultures treated with IL-15 and IL-21 did not induce strong changes in gene expression in the IELs (Supplementary Fig. 6b) or organoids (Supplementary Fig. 6c), further indicating that IFN-γ alone is insufficient to cause cell death under these conditions. In contrast, TNF and FASLG were increased in BIS1 treated co-cultures (Fig. 5e,f), suggesting that BIS1-triggered cytotoxicity may be driven by canonical death-inducing pathways.

Collectively, these data demonstrate that cytokine stimulation enhances GzmB expression and IFN-γ secretion without inducing epithelial apoptosis, whereas enforced IEL–epithelium engagement through BIS1 triggers pronounced cytotoxicity. This establishes the platform we have developed as a sensitive, physiologically relevant model for dissecting immune–epithelial interactions and evaluating modulators of intestinal inflammation and barrier integrity.

## Discussion

We have established an autologous human duodenal IEL–organoid co-culture platform that enables controlled, repeated analysis of lymphoepithelial interactions from minimal biopsy material. By combining expansion and banking of biopsy-derived IELs with a shared co-culture medium that sustains both epithelial and lymphocyte compartments, we have addressed two recurring barriers in human IEL modeling—limited primary cell yield and the incompatibility of immune and epithelial media. In addition, we adapted a Matrigel-bed format ^29^ to improve immune access to the epithelial surface and assay tractability. Using this platform, we show that the co-culture system is capturing both steady-state and perturbed immune–epithelial interactions. Under baseline co-culture conditions, IELs induce an epithelial interferon-associated response, demonstrating that the system supports biologically meaningful bidirectional communication. In addition, exposure to IL-15 and IL-21 and enforced CD3×EpCAM engagement with the bispecific antibody BIS1 ^37,38^ show that the platform can detect enhanced IEL effector activity and epithelial apoptosis. This validates the model’s functional utility and illustrates its potential for studying both homeostatic and injurious IEL– epithelial interactions and for testing potentially harmful compounds.

Recent advances with autologous mucosal T cell–organoid co-cultures have shown that patient lymphocytes can drive epithelial injury in Crohn’s disease ^25^, whereas more complex systems have preserved or reconstituted tissue-resident immune compartments in intestinal organoids ^23,24^. Our model complements these approaches by adopting a more reductionist design centered on only a defined IEL-enriched CD8⁺ T cell compartment, while expansion and banking provide sufficient cell yield for repeated experimentation from minimal biopsy input. However, because this immune compartment is derived from expanded, banked, CD103-enriched CD8⁺ T cells, its phenotype is likely to be partially shaped by *in vitro* culture and TGF-β conditioning. In addition, the expansion protocol may introduce clonal or functional bias. Therefore, we believe these cells should be regarded as IEL-enriched or tissue-resident-like CD8⁺ T cells that retain many functional properties, including the ability to cause cytotoxic cell death through the canonical apoptosis pathway.

The co-culture conditions we developed preserved the biological state of both the epithelial and immune compartments. Organoids retained epithelial morphology and growth, while IELs remained phenotypically and transcriptionally aligned with IELs cultured under IEL-specific conditions. Within this stabilized setting, basal IEL–organoid co-cultures induced an interferon-associated epithelial program, but the IEL compartment remained comparatively stable. This suggests that resident-like IELs can instruct epithelial defense programs without overt cytotoxic activation, even with the relatively high IEL-to-organoid ratios seen in the co-cultures. This is consistent with prior evidence that epithelial IFN-γ sensing can actively shape immune–epithelial crosstalk and exert protective regulatory functions, rather than causing epithelial injury ^41,42^. IFN-γ-exposed intestinal organoids have also been shown to upregulate chemokine programs, in particular CXCL9, CXCL10, and CXCL11, which promote T cell migration without further increasing T cell activation ^43^.

We note that cytokine-driven effector priming could be experimentally uncoupled from epithelial killing in our model. IL-15 and IL-21 increased the levels of effector-associated molecules, including IFN-γ and granzyme B, but did not induce cytotoxic activity of IELs towards organoids. This priming is consistent with the established role of IL-15 in licensing tissue-destructive cytotoxic programs in celiac disease ^3,44^ and with evidence that IL-21 can further augment this IEL cytotoxic potential ^36,40^. However, the absence of measurable epithelial cell death despite increased effector molecule expression suggests that cytokine-driven priming alone is insufficient to trigger cytotoxicity under these conditions. It is likely that additional stress, pro-inflammatory signals, or contact-dependent signals like antigen-dependent T cell receptor−HLA interactions, non-classical HLA-mediated recognition pathways, or activating NK receptor signaling are required to cause cell death and epithelial tissue injury ^3,45^. In line with this, the gene expression of organoids was unchanged upon addition of IL-15 and IL-21 and in co-cultures, indicating that IL-15/IL-21 act only to prime IEL effector functions and do not make epithelial cells susceptible to cell death.

We only observed overt epithelial killing when immune–epithelial engagement was artificially induced using BIS1, a clinical-grade CD3×EpCAM bispecific antibody that promotes antigen-independent CD3/TCR- complex activation ^20,21^. BIS1-induced epithelial death was mediated by strong TNF- and FASLG-associated responses, consistent with activation of cytotoxic effector pathways that may contribute to epithelial apoptosis. The magnitude of this apoptotic response, together with the absence of comparable signals in basal and cytokine-primed conditions, indicates that the co-culture assay can capture robust epithelial killing when an appropriate cytotoxic trigger is present. The absence of apoptosis under IL-15 and IL-21 stimulation thus likely reflects true biology rather than assay insensitivity. More broadly, this demonstrates that the platform can capture epithelial toxicity induced by drugs or bispecific antibodies, in line with previous donor-matched organoid and tumoroid studies of T cell−engaging bispecifics ^46^.

In sum, our co-culture model provides a donor-matched human platform for mechanistic interrogation of lymphoepithelial crosstalk, with expandable and bankable epithelial and immune compartments that make the system scalable for repeated experimentation from minimal biopsy input. It is well suited for modeling disease in contexts such as celiac disease and IBD, where it can be used to define how cytokine cues, epithelial stress signals, and T cell–mediated immune interactions shape epithelial outcomes. In addition, the robust BIS1 response we observed highlights our model’s potential as a preclinical system to evaluate immune-mediated intestinal toxicity, including cytotoxic effects triggered by T cell bispecific antibodies, CAR T cell treatments, and related therapeutic interventions.

## Methods

### Human subjects and biopsy collection

Individuals included in this study consented to be included in the CeD North Netherlands (CeDNN) cohort^47^ and were undergoing endoscopies for duodenal biopsies for symptoms of upper gastrointestinal disorders. The CeDNN study was ethically approved by the medical ethical committee of the University Medical Center Groningen (METc number 2013/440). For this study, biopsies for establishing temporal IEL lines and duodenal organoid lines were included from in total 8 males and 11 females (average age 14.4 ± 4.6 years) (Supplementary Table 16). Only individuals without a celiac disease diagnosis and no further histological evidence of duodenal inflammation were used for functional downstream analyses and co-culture experiments. Biopsies were collected and cryopreserved as previously described ^27^. In brief, two biopsies were collected per 50 ml collection tube in RPMI1640 supplemented with L-glutamine (Gibco) and 10% heat-inactivated fetal bovine serum (FBS). Biopsies were kept on ice until cryopreservation in cold freezing medium (90% heat-inactivated FBS with 10% DMSO) in long-term liquid nitrogen storage.

### IEL isolation

Cryopreserved duodenal biopsies were thawed rapidly per 2 biopsies in a 37 °C water bath (< 60 seconds) and transferred to thawing medium of RPMI 1640 supplemented with 10% FBS. After centrifugation at 300 × g for 5 minutes at room temperature, biopsies were minced on ice into fragments of approximately 1 mm³ and transferred to enzymatic digestion solution containing 200 U/mL collagenase IV (Sigma-Aldrich) and 20 iU/mL DNase II (Sigma-Aldrich) in RPMI 1640 supplemented with 2% FBS. Tissue fragments were digested for 30 minutes at 37 °C under continuous agitation, with intermittent mechanical dissociation by pipetting every 10 minutes. Digestion was terminated by addition of 0.5 M EDTA, followed by centrifugation and a 5-minute incubation in TrypLE Select to further dissociate remaining tissue. Enzymatic activity was then quenched with ice-cold RPMI 1640 containing 10% FBS. Cell suspensions were washed in cold buffer containing DPBS, 0.4% BSA, and 20 iU/mL DNase II, filtered through a 70 μm cell strainer, and pelleted by centrifugation at 300 × g for 5 minutes at 4 °C. Finally, cells were resuspended in FACS buffer (DPBS + 2% FBS), counted by trypan blue exclusion, and left on ice until downstream processing. Following biopsy dissociation into single-cell suspensions, cells were stained for 30 minutes on ice in the dark with antibodies against CD8, CD45, CD4, TCRαβ, TCRγδ, CD103, and CD326/EpCAM. DAPI was used to exclude non-viable cells. After staining, cells were washed, resuspended in FACS buffer, and sorted on a MoFlo Astrios Cell Sorter. IELs were defined as live CD45⁺CD8⁺TCRαβ⁺CD103⁺ cells, while CD326⁺ epithelial cells, dead cells, and non-target lymphocyte populations were excluded. Sorted IELs were recovered in 10% human serum-containing RPMI 1640 medium supplemented with 10% heat-inactivated human serum and IL-2 (200 U/ml) and used for subsequent *in vitro* expansion.

### IEL expansion

Sorted IELs were expanded as previously described ^4,36,48,49^. Briefly, sorted IELs were pelleted by centrifugation at 500 × g for 5 minutes. For primary expansion, cells were plated in round-bottom 96-well plates in IEL medium (RPMI 1640 containing 10% heat-inactivated human serum, IL-2 (200 U/mL), PHA-L (1 μg/mL)), along with irradiated feeder cell mix. The feeder cell mix consisted of pooled peripheral blood mononuclear cells (PBMCs) from three donors obtained from Sanquin and pooled Epstein–Barr virus-transformed B cells (EBV-B cells) from three donors. Both feeder populations were thawed, washed, filtered, and irradiated with 30 Gy (using a IBL 637, Cesium-137 sourced irradiator), counted, and combined prior to use. In the first expansion round, feeder cells were added at a ratio of 200 PBMCs and 20 EBV-B cells per IEL, and IELs were seeded at 5 cells/μL in a total volume of 200 μL per well. Cultures were maintained for 21 days and split 1:2 every 4 days. After 21 days, IELs were harvested and cryopreserved in human serum + 10% DMSO at 7.5 × 10^5^ IELs per cryovial.

For downstream assays, including IEL–organoid co-culture experiments, cryopreserved IELs were thawed and subjected to a second feeder-supported expansion round. Briefly, IELs were washed, counted, and transferred to T25 flasks containing IEL medium (RPMI 1640 supplemented with 10% human serum, IL-2 (100 U/mL), PHA-L (1 μg/mL)) and freshly prepared irradiated feeder cells. In this re-expansion phase, pooled PBMCs and pooled EBV-B cells from three donors each were again used as feeder cells, at a ratio of 100 PBMCs and 10 EBV-B cells per IEL. For each T25 flask, 1.2 × 10^5^ IELs were cultured in 12 mL total volume containing 1.2 × 10^7^ PBMCs and 1.2 × 10^6^ EBV-B cells. Cultures were maintained for 10 days, with 7 mL fresh RPMI 1640 supplemented with IL-2 (100 U/mL) added every other day. To enhance the tissue-resident phenotype indicated by increased expression of CD103, TGF-β (10 ng/mL) was added on day 9, 24 hours prior to downstream assays.

### Generation and maintenance of biopsy-derived duodenal organoids

For organoid establishment, two additional donor-matched cryopreserved duodenal biopsies were thawed and processed separately from biopsies used for IEL isolation. After thawing, tissue was minced into fragments of approximately 1 mm³ with a scalpel and digested in enzymatic digestion solution containing 200 U/mL collagenase IV (Sigma-Aldrich) and 20 iU/mL DNase II (Sigma-Aldrich) in RPMI 1640 supplemented with 2% FBS for 35–40 minutes at 37 °C under continuous agitation. Digestion was stopped by addition of Adv DMEM:F12+++ (Advanced DMEM/F12 supplemented with 100 U/mL penicillin/streptomycin, 1× L-glutamine, and 10 mM HEPES) supplemented with 10% FBS and 10 µM Y-27632 (Rock inhibitor), after which the suspension was washed, filtered through a 100 μm cell strainer, and further washed in ice-cold buffer containing DPBS + 0.4% BSA + 20 iU/mL DNAse II + 10 µM Rock inhibitor. The resulting epithelial fragment─containing pellet was resuspended in 70% Matrigel, seeded as 15 μL domes in pre-warmed 24-well plates, polymerized inverted for 10 minutes at 37 °C, and overlaid with standard organoid medium (Advanced DMEM/F12+++ supplemented with 10% Wnt-conditioned medium (harvested from ATCC L Wnt-3A (CRL-2647) corresponding to 100 ng/ml Wnt3A, (determined by Wnt3A ELISA according to manufacturer’s instructions), R-spondin-1 (100 ng/mL), Noggin (100 ng/mL), B27 (2%), N-acetylcysteine (1.25 mM), nicotinamide (10 mM), EGF (50 ng/mL), SB202190 (10 μM), A83-01 (500 nM), prostaglandin E2 (1 μM), gentamicin (50 μg/mL), and primocin (100 μg/mL), supplemented with 10µM ROCK inhibitor for the first 24 hours. After initial outgrowth, organoids were mechanically disrupted, passaged, and expanded according to established protocols for adult stem cell─derived human intestinal organoids ^14^. Stable organoid lines typically formed within approximately 3 weeks, after which cultures were cryopreserved in CryoStor CS10. For downstream experiments, cryopreserved organoid lines were thawed and re-established in the same standard expansion conditions.

### IEL–organoid co-culture setup

For co-culture experiments, flat-bottom 96-well plates were prepared with 50% Matrigel beds. Plates were placed on ice, and each well was pre-moistened with 200 μL AdvDMEM+++, after which the medium was aspirated without allowing the wells to dry completely. Subsequently, 15 μL of a 1:1 mixture of Matrigel and organoid growth medium supplemented with ROCK inhibitor was added to the center of each well and allowed to polymerize for at least 20 minutes at 37 °C, thereby forming a solid 50% Matrigel bed covering the entire surface of the well ^29^.

For seeding onto Matrigel beds, biopsy-derived organoids maintained in standard expansion culture were dissociated into single cells by disruption of Matrigel domes in ice-cold AdvDMEM+++ and spun down at 450 x g for 5 minutes. Supernatant and residual Matrigel was removed by aspiration, followed by brief incubation in pre-warmed TrypLE with ROCK inhibitor at 37 °C and repeated mechanical resuspension. TrypLE was inactivated with excess ice-cold AdvDMEM+++, after which cells were filtered through a 40 μm strainer, pelleted, and resuspended in organoid growth medium supplemented with 2% Matrigel and ROCK inhibitor. Resuspended organoid cells were seeded directly onto the solidified 50% Matrigel beds at 100 μL per well at a density of 35 cells/μL, corresponding to 3,500 cells per well. Cultures were maintained for 7 days to allow formation of luminal organoids prior to immune co-culture.

For co-culture initiation, day-10 re-expanded IELs were harvested from feeder-supported cultures, counted, pelleted, and resuspended in the indicated assay medium. IEL medium consisted of RPMI 1640 supplemented with 10% human serum and IL-2 (100 U/mL). Co-culture medium consisted of Adv DMEM/F12+++ supplemented with 10% Wnt-conditioned medium, R-spondin-1 (100 ng/mL), Noggin (100 ng/mL), B27 (2%), N-acetylcysteine (1.25 mM), EGF (50 ng/mL), gentamicin (50 μg/mL), primocin (100 μg/mL), IL-2 (100 U/mL), and IL-7 (10 ng/mL). Where indicated, co-cultures were further supplemented with IL-15 (20 ng/mL), IL-21 (3 ng/mL), or BIS1 (200 ng/mL). Before addition of IELs, the original organoid growth medium was removed, and IELs were added at 1 × 10^5^ cells per well.

### Live-cell imaging

For visualization of IEL behavior in co-culture, re-expanded IELs were labeled with CellTracker Deep Red (10 μM) in RPMI 1640 containing 100 U/ml IL-2 for 45 minutes at 37 °C, protected from light. After staining, IELs were pelleted at 300 × g for 5 minutes, resuspended in the appropriate assay medium, and added to pre-established organoid cultures. Time-lapse imaging was performed on a Zeiss Celldiscoverer 7 using brightfield and RFP channels, with images acquired every 20 minutes for 20 hours.

### Organoid surface area measurements

Organoid surface area was quantified using the same seeding strategy described for IEL–organoid co-culture experiments. On day 7, cultures were switched to the indicated media conditions and transferred to an IncuCyte S3 Live-Cell Analysis System (Sartorius), where brightfield images were acquired for indicated time points. Organoid surface area was quantified using the integrated IncuCyte organoid analysis module. For each donor and time point, surface area values were normalized to the corresponding organoid expansion medium condition to calculate relative size differences compared to standard organoid medium.

### Caspase 3/7-based epithelial cell cytotoxicity assay

IEL–organoid co-cultures and corresponding IEL-only and organoid-only control cultures were established as described above. Apoptosis was monitored using the CellEvent Caspase-3/7 Detection Reagent (Invitrogen), which was added directly to the culture medium at the start of the assay, according to the manufacturer’s instructions. Plates were transferred to an IncuCyte S3 Live-Cell Analysis System (Sartorius) and imaged by time-lapse microscopy using phase-contrast and green fluorescence acquisition. Images were acquired in standard scan mode at repeated intervals of 1 hour over the course of the experiment. Caspase-3/7-positive objects were identified in the IncuCyte software using a green object analysis mask. The primary output used for downstream analysis was the raw Total Green Object Integrated Intensity (GCU × μm^2^/image) value exported directly from the software. This integrated intensity metric combines fluorescence intensity and fluorescent object area within the defined green mask. IEL- and organoid-only monocultures served as baseline controls to assess spontaneous apoptosis. Puromycin-treated wells (100 μg/ml) were included as positive controls to induce maximal caspase-3/7 activation. For each biological replicate, the peak caspase signal observed in puromycin-treated wells was defined as the maximal apoptosis (Total Green Object Integrated Intensity (GCU × μm^2^/image) reference and used for normalization of the caspase signal measured across conditions within that biological replicate. This approach allowed comparison of apoptosis kinetics between co-culture conditions while accounting for inter-donor variability in maximal signal intensity.

### Cell collection and preparation for flow cytometric analyses

For flow cytometric analysis of co-cultures, organoid-only cultures, and IEL monocultures, cells were harvested 12 hours after initiation of cultures. Supernatants were removed and cells were incubated with 100 μL Cell Recovery Solution per well for approximately 30−60 minutes at 4 °C to dissolve the Matrigel matrix. Collected cells were dissociated into single cells as described above.

Eventually, single-cell suspensions were pelleted at 400 × g for 5 minutes at 4 °C and stained with 1× Zombie Aqua viability dye for 30 minutes at 4 °C. Unstained control samples received FACS buffer only. After viability staining, cells were washed twice with FACS buffer by centrifugation at 400 × g for 5 minutes. Samples were subsequently fixed in 4% paraformaldehyde for 10 minutes at room temperature, washed once more with FACS buffer, and finally resuspended in FACS buffer for storage at 4 °C until antibody staining and flow cytometric analyses.

### Flow Cytometry analyses

For medium-compatibility assays, IELs were cultured for 24 hours in the indicated media conditions, and viability was assessed with propidium iodide on a Novocyte Quanteon flow cytometer (Agilent) and processed with NovoExpress software (Agilent).

For phenotypic flow cytometry, harvested cells were stained with Zombie Aqua prior to fixation, as described above. Antibodies against CD8, CD103, PD-1, TIGIT, GzmB, CCR9, CD57, CD326/EpCAM, and CD45RA were used for phenotypic flow cytometry. All stainings were performed using 1× BD Perm/Wash buffer. Cells were permeabilized for 15 minutes at room temperature in the dark, stained for 30 minutes on ice in the dark, washed twice in BD Perm/Wash buffer, and resuspended in FACS buffer. Data were acquired on a Cytek Aurora spectral analyzer and processed with SpectroFlow software. IELs were analyzed by gating on non-debris, singlet, live CD8⁺EpCAM⁻CD45RA⁻ cells (Supplementary fig. 1B).

### Olink analysis of conditioned media

Conditioned media were harvested 12 hours after culture initiation, centrifuged at 1000 × g for 5 minutes at 4 °C, and transferred to fresh plates to remove residual cell material. Conditioned media were stored at −80 °C until analysis by SciLifeLab Affinity Proteomics (Uppsala, Sweden) using the Olink Target 96 Immuno-Oncology panel. Output data were provided as NPX values on a log2 scale, relative to Olink proprietary internal standards.

Olink data were processed and filtered in R (version 4.4.2) prior to downstream analysis. For each assay, blank-corrected delta NPX values (d_NPX) were calculated by subtracting the NPX value of the corresponding condition-matched blank control from each measurement. Assays with high background signal in blank samples were excluded using a threshold of blank NPX > 6. After blank-based filtering, data were subset to the conditions and sample types relevant to each analysis. Statistical testing was then performed on d_NPX values using paired, two-sided t-tests on donor-matched samples, followed by Benjamini-Hochberg correction for multiple testing. Proteins were considered significantly differentially abundant when at least one pairwise comparison reached significance after Benjamini-Hochberg correction (adjusted p-value < 0.05).

### ELISA of conditioned media

Wnt-conditioned medium for organoid culture was harvested from ATCC L Wnt-3A cells (CRL-2647) and ran over a 0.2 μm filter (Thermo Scientific, 568-0020) to remove residual cell material. Conditioned media from other cultures were collected 12 hours after culture initiation, centrifuged at 1,000 × g for 5 minutes at 4 °C, and transferred to fresh plates to remove residual cell material. Conditioned media were stored at −80 °C until analysis. Wnt3A, granzyme B, and IFN-γ concentrations were determined by ELISA using RCD Systems DuoSet kits according to the manufacturer’s instructions (Wnt3A: DY1324B-05; granzyme B: DY2906-05; IFN-γ: DY285B-05).

### RNA isolation from mono- and co-cultured IEL and epithelial compartments

For RNA-seq analysis, samples were harvested 6 hours after initiation of culture. For monoculture control conditions, IELs and organoids were lysed directly without further separation. IELs were collected in ice-cold PBS, pelleted by centrifugation at 400 × g for 5 minutes, and lysed in RLT buffer. For organoid-only conditions, culture medium was removed and organoids were lysed directly in the wells by addition of RLT buffer. Lysates were collected and stored at −80 °C until RNA isolation.

For co-culture samples, the epithelial and immune compartments were separated prior to RNA isolation. Residual supernatant was removed, and 100 μL Cell Recovery Solution was added to each well to dissolve the Matrigel bed. Plates were incubated at 4 °C for 30–60 minutes, after which suspensions from replicate wells of the same condition were pooled and transferred to 2 mL tubes. Samples were centrifuged at 400 × g for 5 minutes and resuspended in EasySep buffer (Stem Cell Technologies). To deplete IELs from epithelial-containing suspensions, 50 μL a-CD3 antibody selection cocktail (Stem Cell Technologies) was added and incubated for 5 minutes at room temperature, followed by addition of 50 μL magnetic beads (Stem Cell Technologies) and a further 5-minute incubation. Tubes were placed in a magnet for 10 minutes to retain labeled cells. The unlabeled supernatant was transferred to fresh tubes and subjected to a second magnetic depletion step. The pooled unlabeled fraction was collected as the epithelial compartment. The magnet-retained fraction was collected as the IEL compartment after additional washes in EasySep buffer (Stem Cell Technologies). Both fractions were pelleted by centrifugation at 400 × g for 5 minutes and lysed in RLT buffer. Lysates were stored at −80 °C until RNA extraction. RNA was isolated using the Qiagen RNeasy MinElute protocol according to the manufacturer’s instructions.

### RNA-sequencing and data pre-processing

RNA samples isolated from organoid, IEL, and separated co-culture compartments were used for bulk RNA-sequencing. Libraries were prepared using a eukaryotic strand-specific mRNA library preparation workflow with reference genome alignment for human samples. Sequencing was performed on an Illumina NovaSeq platform using paired-end 150 bp reads, with a target output of 6 Gb per sample. Raw sequencing data was quality controlled using FastQC (1). Alignment and quantification of transcripts was done using Salmon v1.9^50^ and the reference human genome hg38 (salmon_sa_index) downloaded from Refgenie^51^. MultiQC^52^ was used to aggregate the quality reports from the different tools and to assess overall quality of sequencing and alignment. Transcripts counts were imported into R (v4.4.1) and converted to a gene counts matrix using tximport ^53^ and Ensembl (v113) to map transcripts to genes. We first performed an exploratory analysis of the data using edgeR^54^ to carry out principal component analysis, which identified that RNA integrity and GC content represented unwanted sources of variation that should be corrected in downstream analysis, as well as identifying cross-contamination between cell types.

### Estimation and correction of contamination

To correct for cross-contamination in the co-culture samples, we used hspe ^55^, a deconvolution method that uses the gene expression of marker genes to estimate the proportion of each cell type in a mixed sample. To define the marker genes for each cell type, we ran a differential expression test using edgeR to find genes differentially expressed between pure organoids and pure IELs per experimental condition. We then selected the top 100 DEGs with a log2FC > 5 and an adjusted p-value < 0.0001 as marker genes. We validated the accuracy of the model for predicting cell type proportions with these marker genes by testing the model in artificial mixed samples made from pure organoids and IEL samples in known proportions. Finally, we used the validated model to estimate the proportion of contaminating transcripts from IELs present in co-culture organoid samples, and vice-versa.

### Differential expression and pathway enrichment analyses

Testing for DEGs was done using edgeR and limma-voom ^56^, filtering out lowly expressed genes (> 500 counts) and including donor as a random effect and RNA integrity and GC content as fixed effects in the model. The Benjamini-Hochberg method was used for multiple testing correction, and genes with |log2FC| > 1 and an adjusted p-value < 0.05 were considered differentially expressed.

To find the DEGs in comparisons between co-culture and pure samples that are truly an effect of the co-culture and not a result of contamination, we used a multi-step filtering strategy. First, from the co-culture sample, we filtered out genes that were highly expressed exclusively in the pure sample of the contaminating cell type (in each experimental condition). We then used the pure samples to create artificial mixed samples with the same proportion of IEL/Organoid transcripts, estimated as described above in “Estimation and correction of contamination”. DEGs found in this step were designated “contaminating DEGs”. Finally, we ran the test for the real co-culture samples versus pure samples and filtered out the “contaminating DEGs” from the results. Gene set enrichment analysis and over-representation analysis were done using the package clusterProfileR ^57^ and the databases Gene Ontology Biological Process, KEGG Pathway, and Reactome. Enriched terms with adjusted p-value < 0.05 were kept.

### Statistical analyses

Data are presented as donor-matched measurements. Boxplots show the median and interquartile range (IQR: 25^th^–75^th^ percentile), with individual donor values overlaid as points. Whiskers represent values within 1.5 × IQR. Time-course data are shown as median with IQR across donors. For time-course experiments, the area under the curve (AUC) was calculated per donor using the trapezoidal rule. Statistical comparisons between paired conditions were performed using paired t-tests. For multiple comparisons, p-values were adjusted using Benjamini-Hochberg correction. Paired Wilcox test with Benjamini-Hochberg correction was used to assess significance levels in non-parametric data. Significance levels were set at *P < 0.05, **P < 0.01, ***P < 0.001, and ****P < 0.0001. Statistical analysis was performed and graphs were generated using R-studio (R version 4.4.2).

## Data availability

RNA sequencing data will be made available in the GEO database. Lists of Olink assays and of DEGs of the analyses described this study are provided in the Supplementary Information, Supplementary tables and Source Data file. Source data are provided with this paper.

**Table 1.** Core media, buffers, and matrices.

| Reagent/material | Source | Cat. No. | Working concentration |
| --- | --- | --- | --- |
| Advanced DMEM/F12 | Thermo Fisher Scientific | 12634010 | NA |
| RPMI 1640 | Thermo Fisher Scientific | 11875093 | NA |
| FBS (fetal bovine serum) | Thermo Fisher Scientific | A5256701 | 10% |
| DMSO | Sigma-Aldrich / Merck | D2650-100ML | 10% |
| Human serum | Sigma-Aldrich | H4522-100ML | 10% |
| Penicillin/Streptomycin | Thermo Fisher Scientific | 15140122 | 100 U/mL |
| L-glutamine (100x) | Thermo Fisher Scientific | 25030081 | 1x |
| HEPES | Thermo Fisher Scientific | 15630056 | 10 mM |
| DPBS | Thermo Fisher Scientific | 14190-169 | NA |
| MACS® BSA Stock Solution | Miltenyi Biotec | 130-091-376 | 0.4% |
| Matrigel | Corning | 354234 | 70% & 50% |
| Wnt-conditioned medium | ATCC L Wnt-3A cells | CRL-2647 | 10% (= 100 ng/ml Wnt3A) |

**Table 2.**
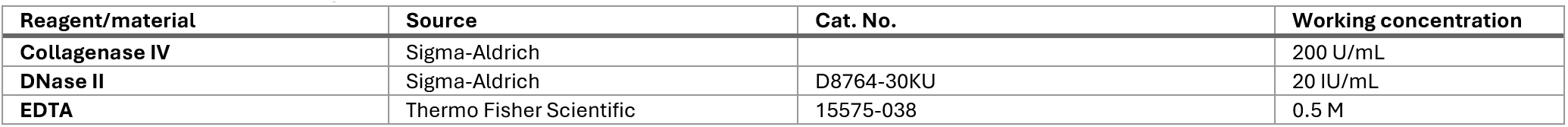
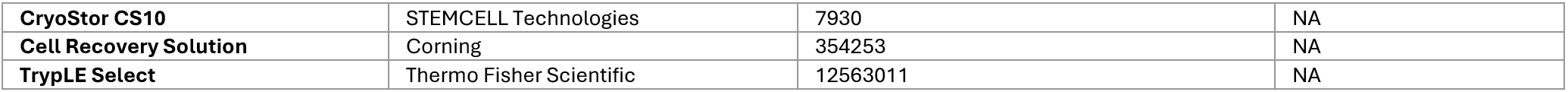
Cell- and tissue processing materials.

| Reagent/material | Source | Cat. No. | Working concentration |
| --- | --- | --- | --- |
| Collagenase IV | Sigma-Aldrich |  | 200 U/mL |
| DNase II | Sigma-Aldrich | D8764-30KU | 20 IU/mL |
| EDTA | Thermo Fisher Scientific | 15575-038 | 0.5 M |
| **CryoStor CS10** | STEMCELL Technologies | 7930 | NA |
| **Cell Recovery Solution** | Corning | 354253 | NA |
| **TrypLE Select** | Thermo Fisher Scientific | 12563011 | NA |

**Table 3.** Cytokines, growth factors, inhibitors, antibiotics and organoid additives.

| Reagent/material | Source | Cat. No. | Working concentration |
| --- | --- | --- | --- |
| **IL-2** | R&D Systems | 202-IL-010/CF | 200 U/mL & 100 U/mL |
| **PHA-L** | Thermo Fisher Scientific | 00-4977-93 | 1 µg/mL |
| **TGF-β** | R&D Systems | 240-B-010/CF | 10 ng/mL |
| **R-spondin-1** | R&D Systems | 4645-RS-100/CF | 100 ng/mL |
| **Noggin** | R&D Systems | 6057-NG-100/CF | 100 ng/mL |
| **B27 supplement** | Thermo Fisher Scientific | 17504044 | 2% |
| **N-acetylcysteine** | Sigma-Aldrich | A9165-25G | 1.25 mM |
| **Nicotinamide** | Sigma-Aldrich | N0636-100G | 10 mM |
| **EGF** | R&D Systems | 236-EG-200 | 50 ng/mL |
| **SB202190** | R&D Systems | 1264/10 | 10 µM |
| **A83-01** | R&D Systems | 2939/10 | 500 nM |
| **Prostaglandin E2 (PGE2)** | R&D Systems | 2296/10 | 1 µM |
| **Gentamicin** | Thermo Fisher Scientific | 15710064 | 50 µg/mL |
| **Primocin** | InvivoGen | ant-pm-2 | 100 µg/mL |
| **ROCK inhibitor / Y-27632** | R&D Systems | 1254/10 | 10 µM |
| **IL-7** | R&D Systems | 207-IL-005/CF | 10 ng/mL |
| **IL-15** | BioLegend | 570304 | 20 ng/mL |
| **IL-21** | BioLegend | 571204 | 3 ng/mL |
| **BIS1** | Kindly provided by the lab of Wijnand Helfrich | NA | 200 ng/mL |

**Table 4.**
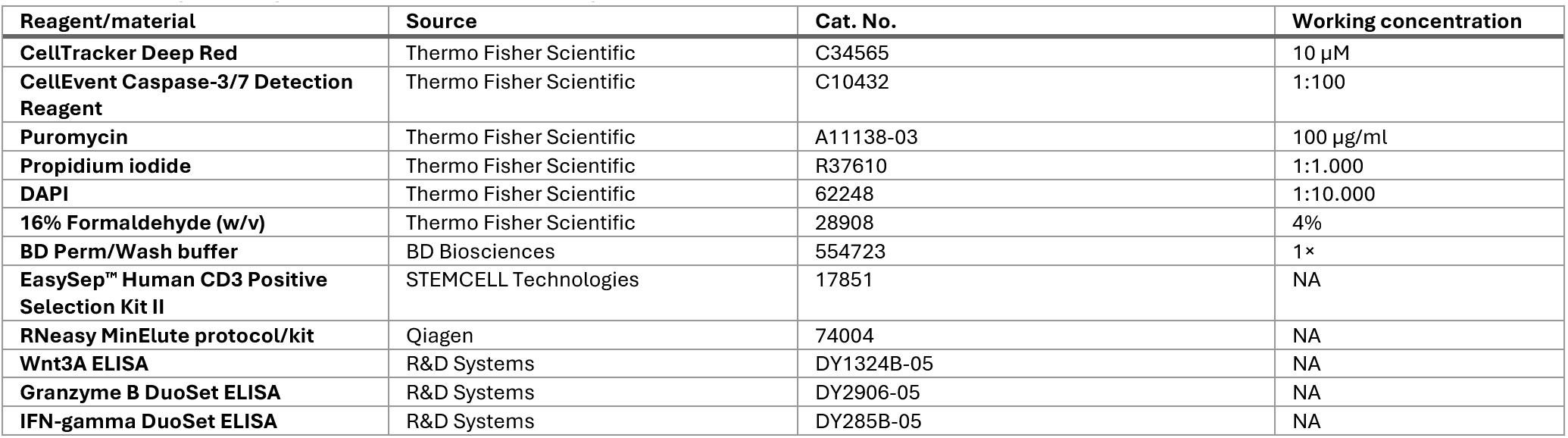
Assay, flow-processing and RNA-separation reagents.

| Reagent/material | Source | Cat. No. | Working concentration |
| --- | --- | --- | --- |
| **CellTracker Deep Red** | Thermo Fisher Scientific | C34565 | 10 µM |
| **CellEvent Caspase-3/7 Detection Reagent** | Thermo Fisher Scientific | C10432 | 1:100 |
| **Puromycin** | Thermo Fisher Scientific | A11138-03 | 100 µg/ml |
| **Propidium iodide** | Thermo Fisher Scientific | R37610 | 1:1.000 |
| **DAPI** | Thermo Fisher Scientific | 62248 | 1:10.000 |
| **16% Formaldehyde (w/v)** | Thermo Fisher Scientific | 28908 | 4% |
| **BD Perm/Wash buffer** | BD Biosciences | 554723 | 1× |
| **EasySep™ Human CD3 Positive Selection Kit II** | STEMCELL Technologies | 17851 | NA |
| **RNeasy MinElute protocol/kits** | Qiagen | 74004 | NA |
| **Wnt3A ELISA** | R&D Systems | DY1324B-05 | NA |
| **Granzyme B DuoSet ELISA** | R&D Systems | DY2906-05 | NA |
| **IFN-gamma DuoSet ELISA** | R&D Systems | DY285B-05 | NA |

**Table 5.** Antibodies for IEL sorting.

| Target | Fluorophore/conjugate | Clone | Source | Cat. No. | Working concentration |
| --- | --- | --- | --- | --- | --- |
| **CD8** | PE | SK1 | BioLegend | 344705 | 1:80 |
| **CD45** | Alexa Fluor 700 | HI30 | BioLegend | 304023 | 1:100 |
| **CD4** | BB700 | SK3 | BD Biosciences | 566393 | 1:80 |
| **TCRαβ** | BV421 | IHC26 | BioLegend | 210120 | 1:80 |
| **TCRγδ** | PE-Cy7 | 11F2 | BD Biosciences | 655410 | 1:20 |
| **CD103** | BB515 | Ber-ACT8 | BD Biosciences | 564578 | 1:40 |
| **CD326/EpCAM** | APC | CO17-1A | BioLegend | 369809 | 1:40 |

**Table 6.** Phenotypic flow cytometry panel.

| Target/reagent | Fluorophore/conjugate | Clone | Source | Cat. No. | Working concentration |
| --- | --- | --- | --- | --- | --- |
| **Zombie Aqua** | Viability dye | NA | BioLegend | 423102 | 1:1000 |
| **CD8** | Spark UV 387 | SK1 | BioLegend | 344775 | 1:80 |
| **CD103** | BV605 | Ber-ACT8 | BioLegend | 350217 | 1:40 |
| **PD-1** | PE-Fire 700 | A17188B | BioLegend | 621621 | 1:50 |
| **TIGIT** | BV421 | A15153G | BioLegend | 372709 | 1:50 |
| **GzmB** | PE | GB11 | Thermo Fisher Scientific | 12-8899-41 | 1:25 |
| **CCR9** | APC-Fire 750 | L053E8 | BioLegend | 358927 | 1:66.67 |
| **CD57** | BB515 | NK-1 | BD Biosciences | 565945 | 1:50 |
| CD326/EpCAM | BUV563 | EBA-1 | BD Biosciences OptiBuild | 748398 | 1:40 |
| CD45RA | PerCP | HI100 | BioLegend | 304155 | 1:50 |

## Acknowledgements

We thank K. McIntyre for critically reading and editing the manuscript. We thank J. Teunis, T. Bijma, G. Mesander for flow cytometry assistance and K. Sjollema for live cell imaging assistance. We thank participants of the Celiac Disease Northern Netherlands cohort that donated duodenal biopsies for supporting research projects with human material. We thank Prof. W. Helfrich for kindly providing BIS1 antibody.

## Author contributions

CW, SW and IHJ conceived the study. JM, CW, SW and IHJ designed the study. JM, SA, JGA, ADRS, HS, ES, RM and RAMM performed experiments. JM, SA and NR analyzed the data. JM, SA, NR, SW and IHJ interpreted the data and wrote the manuscript. GG and MW included human subjects and collected biomaterials

## Correspondence

Correspondence and request for materials should be addressed to Sebo Withoff or Iris Helene Jonkers

