## Supplementary Figures for "Autologous biopsy-derived co-culture platform for interrogation of intestinal epithelial-T cell crosstalk"

### Supplementary Figure 1

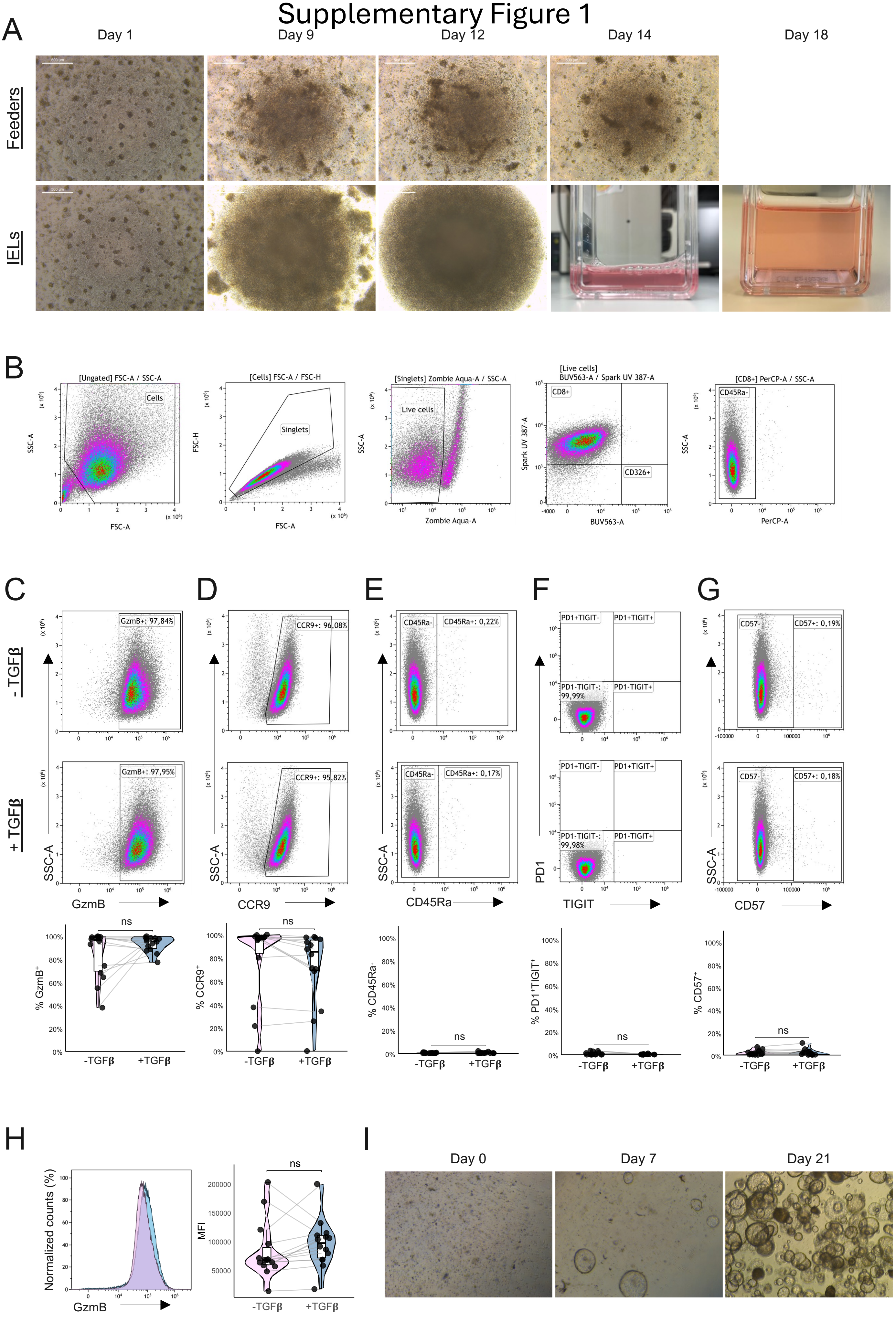

#### Supplementary fig. 1| Phenotypic characterization of expanded IELs and establishment of biopsy-derived organoid lines

**a** Representative images showing feeder-supported expansion of IELs during the second expansion round. **b** Gating strategy used to define live $CD8^{+}EpCAM^{-}CD45RA^{-}$ immune cells for downstream phenotypic analyses. **c–g** Representative flow cytometry plots and quantification of $GzmB^{+}$, $CCR9^{+}$, $CD45RA^{+}$, PD-$1^{+}TIGIT^{+}$, and $CD57^{+}$ IELs after culture with or without TGF-β stimulation (10 ng/mL for 24 h). **h** Quantification of intracellular GzmB levels in $GzmB^{+}$ IELs, shown as median fluorescence intensity (MFI). **i** Representative images showing establishment and expansion of biopsy-derived intestinal organoid lines. In (**c–h**), each dot represents one IEL line (n = 14). Statistical significance was assessed using paired t-tests with Benjamini-Hochberg correction for multiple comparisons where indicated. *P < 0.05, **P < 0.01, ***P < 0.001, ****P < 0.0001.

### Supplementary Figure 2

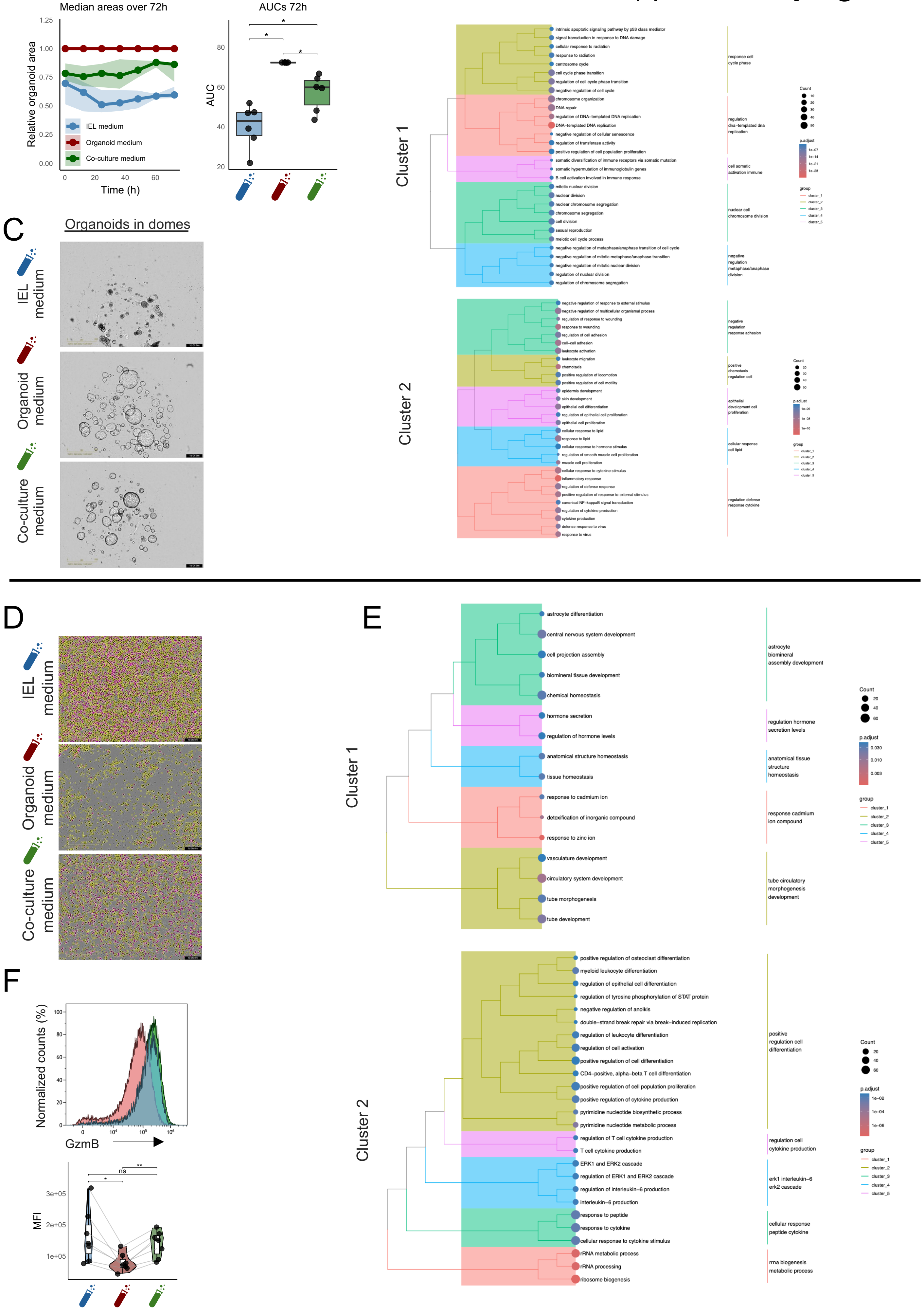

#### Supplementary fig. 2 | Co-culture medium preserves organoid integrity and IEL phenotype over time

**a** Quantification of relative organoid surface area over the 72 h after switching to IEL medium, organoid medium, or co-culture medium. Values were normalized to organoids maintained in organoid medium, which was set to 1. Data in time-course plot represents median ± IQRs. The corresponding area under the curve (AUC) values are shown at right (n = 6 organoid lines). **b** Over-representation analysis of the organoid gene clusters shown in Fig. 2d, with individual pathways displayed separately. **c** Representative brightfield images of organoids grown in Matrigel domes for 24 h in the indicated media. **d** Representative phase-contrast images of IELs 24 h after switching to the media indicated, with IncuCyte-based eccentricity classification shown for low-eccentricity cells (yellow) and high-eccentricity cells (purple). **e** Over-representation analysis of the IEL gene clusters shown in Fig. 2j, with individual pathways displayed separately. **f** Quantification of intracellular GzmB levels in IELs 12 h after switching to the media indicated, shown as median fluorescence intensity (MFI, n = 7 IEL lines). Statistical significance was assessed using paired t-tests with Benjamini-Hochberg correction for multiple comparisons where indicated. $^{*}P < 0.05$, $^{**}P < 0.01$, $^{***}P < 0.001$, $^{****}P < 0.0001$.

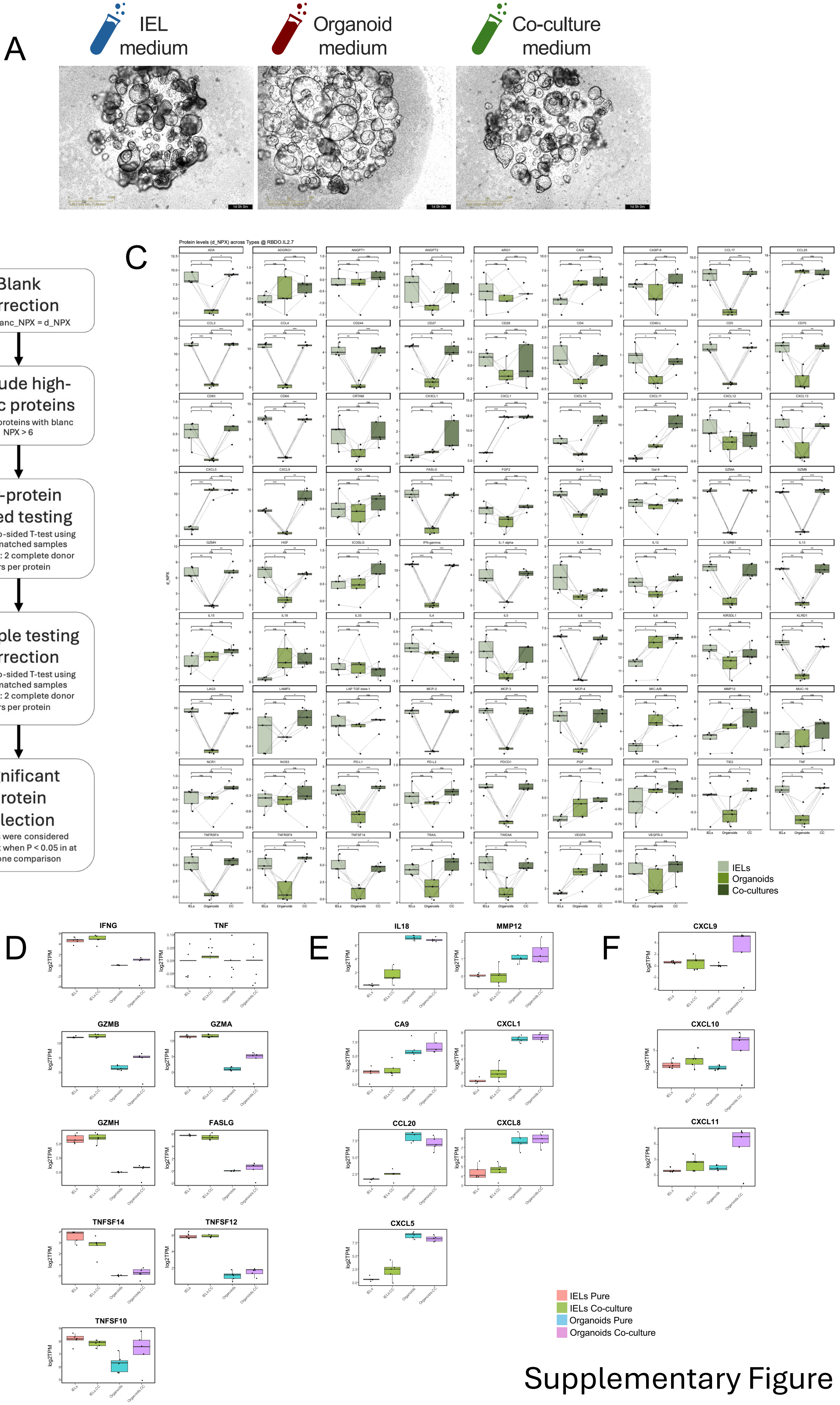


Supplementary Figure 3

#### Supplementary fig. 3 | Compartment-resolved protein and RNA profiles define IEL- and organoid-derived signals in co-culture

**a** Representative brightfield images of intestinal organoids grown in Matrigel domes after addition of IELs and switching to IEL medium, organoid medium, or co-culture medium. **b** Schematic overview of Olink data processing. **c** Individual Olink protein measurements after processing according to (**b**). Conditioned media were collected from IEL monocultures, organoid monocultures, and IEL–organoid co-cultures grown in co-culture medium (n = 5). Values are shown as blank-corrected NPX values (d_NPX). **d**–**f** Expression levels in log2TPMs of selected proteins presented in Figure 3C−E determined by bulk RNA-seq after 6 h (n = 5). For Olink quantifications, each dot represents one conditioned media sample. For Log2TPM plots, each dot represents one donor-matched IEL or organoid line (n = 5). Statistical significance was assessed using paired tests with Benjamini-Hochberg correction for multiple comparisons where indicated. *P < 0.05, **P < 0.01, ***P < 0.001, ****P < 0.0001.

### Supplementary Figure 4

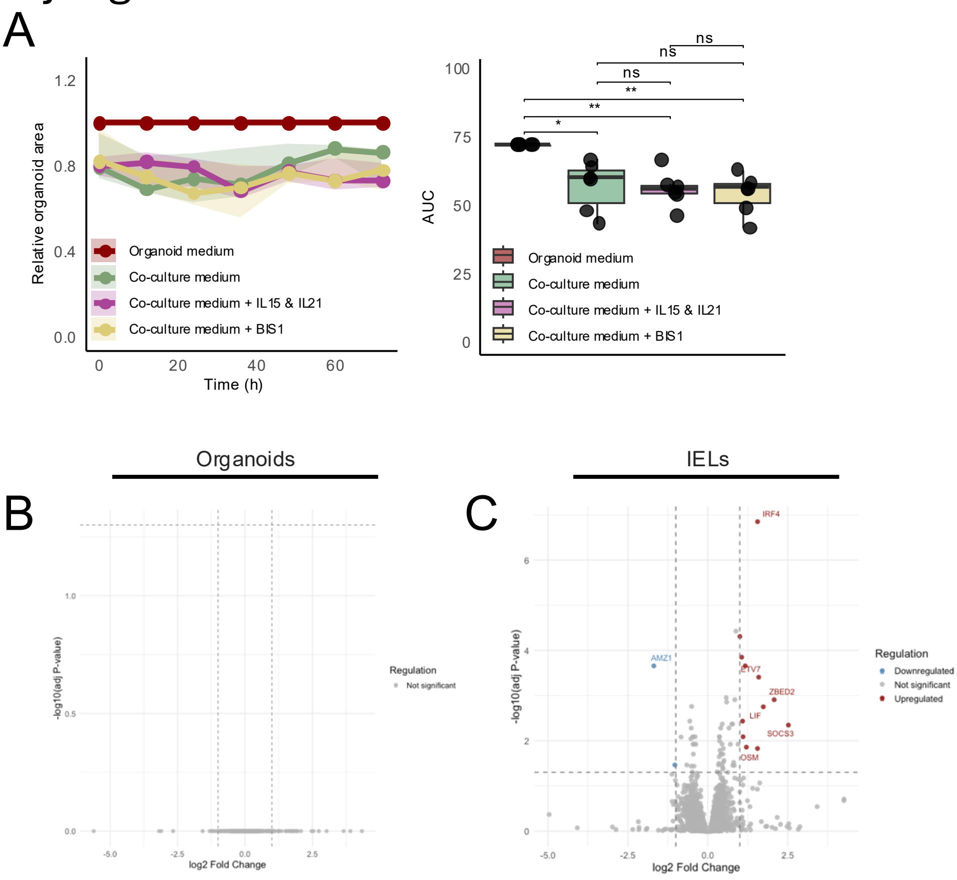

#### Supplementary fig. 4 | IL-15, IL-21 and BIS1 have limited effects on organoid growth and monoculture transcriptomes

**a** Quantification of relative organoid surface area over 72 h after switching to organoid medium, co-culture medium, co-culture medium supplemented with IL-15 and IL-21, or co-culture medium supplemented with BIS1. Values were normalized to organoids maintained in organoid medium, which was set to 1. Data in the time-course plot represent median ± IQRs. Corresponding area under the curve (AUC) values are shown at right (n = 6 organoid lines). **b, c** Volcano plots showing differential gene expression after culture in co-culture medium supplemented with IL-15 and IL-21 or BIS1 compared with unstimulated co-culture medium controls, as determined by bulk RNA-seq for pure (**b**) organoids and (**c**) IELs. Statistical significance was assessed using paired t-tests with Benjamini-Hochberg correction for multiple comparisons where indicated. *$P < 0.05$, **$P < 0.01$, ***$P < 0.001$, ****$P < 0.0001$.

### Supplementary Figure 5

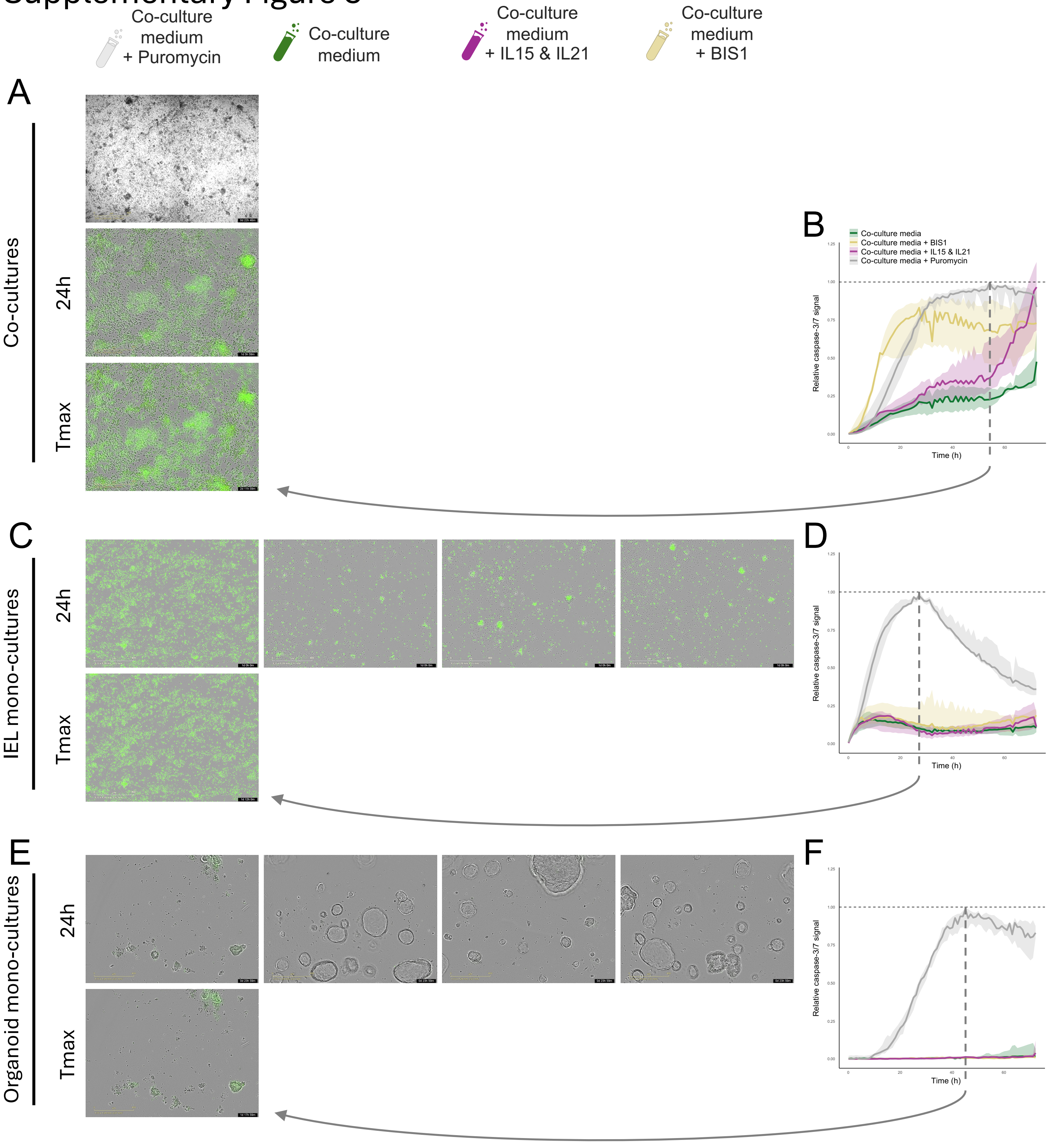

#### Supplementary fig. 5 | Caspase-3/7 controls confirm selective BIS1-induced apoptosis in IEL–organoid co-cultures only

**a** Representative bright field and phase-contrast with caspase-3/7 fluorescence overlay images of IEL–organoid co-cultures treated with puromycin, co-culture medium, co-culture medium supplemented with IL-15 and IL-21, or co-culture medium supplemented with BIS1. Images are shown at 24 h and at the time point of maximal puromycin-induced caspase-3/7 signal (Tmax). **b** Quantification of caspase-3/7 signal over time in IEL–organoid co-cultures under the indicated conditions. For each donor, the maximal caspase-3/7 signal observed in puromycin-treated co-cultures was used as the reference value and set to 1. Data represent medians ± IQRs (n = 7). **c** Representative phase-contrast and caspase-3/7 fluorescence overlay images of IEL monocultures treated under the same conditions, shown at 24 h and Tmax. **d** Quantification of relative caspase-3/7 signal over time in IEL monocultures. Data represent medians ± IQRs (n = 7). **e** Representative phase-contrast and caspase-3/7 fluorescence overlay images of organoid monocultures treated under the same conditions, shown at 24 h and Tmax. **f** Quantification of relative caspase-3/7 signal over time in organoid monocultures. Data represent medians ± IQRs (n = 7). Dashed horizontal lines indicate the normalized maximal puromycin-induced caspase-3/7 signal. Dashed vertical lines indicate Tmax.

### Supplementary Figure 6

A

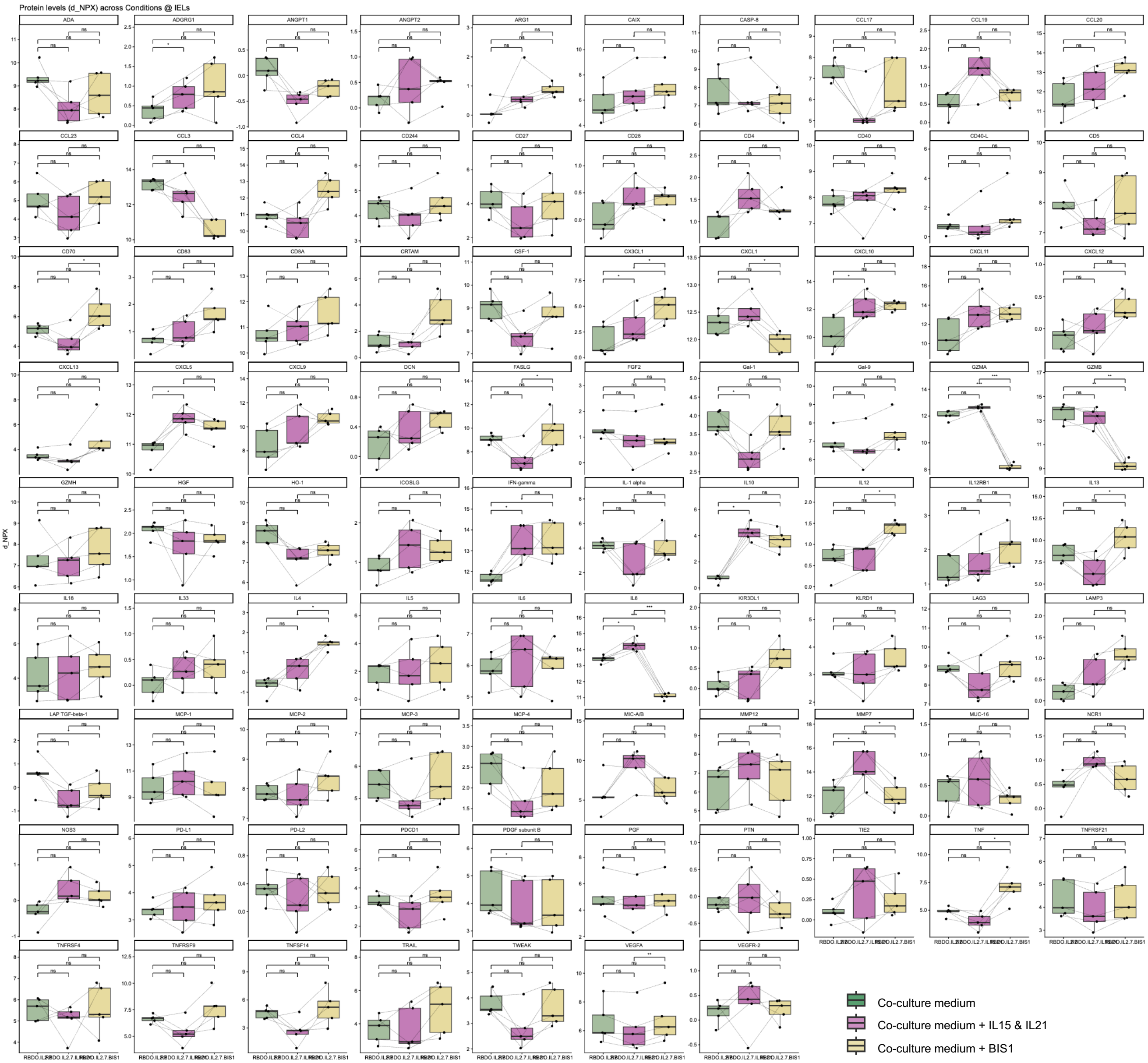


B

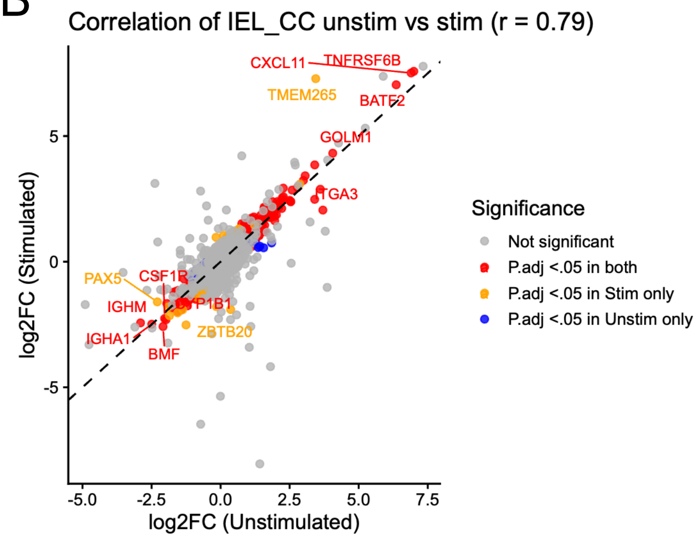


C

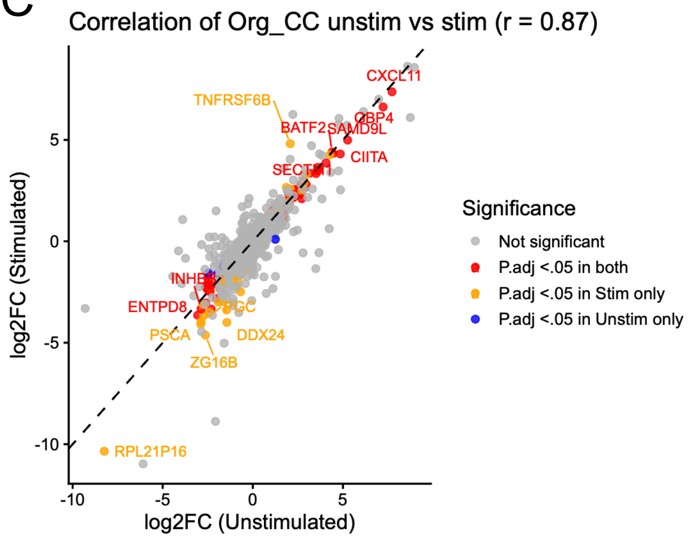

#### Supplementary fig. 6 | Cytokine priming preserves baseline co-culture transcriptional responses, whereas BIS1 induces distinct soluble mediators

**a** Olink measurements of cytokines, chemokines, and immune-mediators in conditioned media from IEL–organoid co-cultures grown in co-culture medium, co-culture medium supplemented with IL-15 and IL-21, or co-culture medium supplemented with BIS1. Values are shown as blank-corrected (d_NPX) values. Each dot represents one conditioned media sample. Statistical significance was assessed using paired t-tests with Benjamini-Hochberg correction for multiple comparisons, where indicated. *P < 0.05, **P < 0.01, ***P < 0.001, ****P < 0.0001. **b** Correlation of RNA-seq log2 fold changes in IELs comparing monocultures and co-cultures under standard co-culture medium conditions versus co-culture medium supplemented with IL-15 and IL-21. Each dot represents one gene. X-axis shows the log2 fold change between IEL monocultures and IELs recovered from co-cultures in standard co-culture medium. Y-axis shows the corresponding log2 fold change in co-culture medium supplemented with IL-15 and IL-21. Genes are colored according to adjusted significance in one or both comparisons. **c** Correlation of RNA-seq log2 fold changes in organoids comparing monocultures and co-cultures under standard co-culture medium conditions versus co-culture medium supplemented with IL-15 and IL-21. Each dot represents one gene. X-axis shows the log2 fold change between organoid monocultures and organoids recovered from co-cultures in standard co-culture medium. Y-axis shows the corresponding log2 fold change in co-culture medium supplemented with IL-15 and IL-21. Genes are colored according to adjusted significance in one or both comparisons. Dashed diagonal lines indicate equal log2 fold change between the two medium conditions.
