## Supplementary Videos Legends for "Autologous biopsy-derived co-culture platform for interrogation of intestinal epithelial-T cell crosstalk"

### Supplementary video 1 | Co-culture medium preserves intestinal organoid morphology

Representative time-lapse videos of intestinal organoids grown in Matrigel domes after switching to IEL medium, organoid medium, or co-culture medium. Organoids were followed for 24 h after medium switch.

### Supplementary video 2 | Matrigel domes restrict IEL access to intestinal organoids

Representative time-lapse videos of intestinal organoids grown in Matrigel domes after switching to IEL medium, organoid medium, or co-culture medium. IELs were added separately to the cultures at the time of medium switch. Cultures were followed for 24 h.

### Supplementary video 3 | IELs remain motile and contact organoids in co-culture medium

Representative time-lapse video of IEL–organoid co-cultures followed for 20 h. Re-expanded IELs were pre-stained with CellTracker Deep Red and added to pre-established organoids. Images were acquired every 20 min using brightfield and red fluorescence channels.
